# Oral and gut microbiomes reveal potential physiological constraints on release readiness in rehabilitating Javan slow lorises

**DOI:** 10.64898/2026.04.23.720515

**Authors:** Abdullah Langgeng, Marie Sigaud, Wendi Prameswari, Nur Purba Priambada, Puji Rianti, Richard Moore, Karmele Llano Sanchez, Wanyi Lee, Andrew J. J. MacIntosh, Ikki Matsuda

## Abstract

Illegal wildlife trade and habitat degradation displace thousands of animals annually in Southeast Asia, with many confiscated primates housed in rehabilitation centers that increasingly function as long-term holding environments. In slow lorises, dental clipping associated with the pet trade may generate persistent disruption along the oral–gut axis, potentially undermining physiological readiness for release in ways not captured by conventional screening. Here, we evaluated whether microbiome structure provides an integrative marker of release readiness in rehabilitating Javan slow lorises (*Nycticebus javanicus*). From June to October 2024, we collected fecal (n = 26) and saliva (n = 18) samples from 19 adults housed at YIARI, including 10 release candidates and 9 non-candidates classified primarily based on tooth loss, medical history, and possibility of release. Bacterial communities were characterized using 16S rRNA (V3–V4) amplicon sequencing, with alpha and beta diversity, taxonomic enrichment (LEfSe), and predicted functional profiles (PICRUSt2) assessed. Microbiome composition was strongly compartmentalized by body site, with higher alpha diversity in the gut. Release candidacy was associated with modest gut compositional differences, whereas oral microbiomes showed pronounced divergence between candidates and non-candidates. Non-candidates were enriched in dysbiosis-associated taxa and degradation-oriented functional pathways, while candidates showed enrichment of biosynthetic and central energy metabolism pathways. Gut microbiome structure was stable across pre-release and soft-release phases. These findings indicate that oral and gut microbiomes represent distinct physiological niches and that persistent oral microbiome alteration is a sensitive marker of long-term dental perturbation. Integrating microbiome-informed metrics may improve multidimensional assessment of release readiness.

## INTRODUCTION

Across Southeast Asia, thousands of wild animals are displaced each year as a result of illegal wildlife trade and ongoing habitat degradation, entering rescue and rehabilitation systems that now function as long-term ecological holding environments rather than short-term medical facilities (Cheyne, 2006; Nijman, 2009). For many confiscated primates, rehabilitation centers represent the first prolonged interface between wild-origin individuals and captive conditions, where veterinary intervention, captive diets, and behavioral management collectively shape subsequent survival trajectories (EAZA, 2022; Moore, Wihermanto, & Nekaris, 2014). In Indonesia, slow lorises comprise a substantial proportion of confiscated primates, and individuals frequently arrive with severe injuries associated with capture, transport, and prior captivity (Fuller, Eggen, Wirdateti, & Nekaris, 2018; Guno, Kusuma, & Taqwim, 2022; Moore et al., 2014). Consequently, rehabilitation has become not only a welfare intervention but also a critical ecological filter influencing population recovery.

Although release protocols have improved substantially, survival following reintroduction remains poorly documented. Post-release mortality is commonly attributed to behavioral maladaptation, chronic stress, and latent health impairment, suggesting that conventional screening, which is typically based on external condition, parasite load, and overt disease, fails to capture integrative physiological readiness for autonomous life in the wild (IUCN-SSC, 2019; Kenyon et al., 2014; van der Sandt, 2017). Such screening frameworks may overlook cumulative physiological changes that develop during captivity and only become apparent after release.

Host-associated microbiomes are now recognized not merely as ancillary components of host physiology, but as integral ecological systems that mediate nutrient assimilation, immune competence, and resilience to environmental stress. In mammals, gut microbial communities regulate energy harvest, fiber fermentation, and short-chain fatty acid production, thereby shaping metabolic efficiency and buffering hosts against nutritional limitation (Amato et al., 2013; Flint, Scott, Duncan, Louis, & Forano, 2012; Mukherjee, Lordan, Ross, & Cotter, 2020). Beyond digestion, the gut microbiome plays a central role in maintaining immune homeostasis, including the regulation of mucosal tolerance, inflammatory tone, and resistance to pathogen colonization (Kamada, Seo, Chen, & Nunez, 2013; Pickard, Zeng, Caruso, & Nunez, 2017). Disruption of these communities, commonly referred to as dysbiosis, describes compositional and functional restructuring of host-associated microbial communities that may alter ecological stability or host–microbe interactions and has been linked to systemic inflammation and metabolic impairment (Carding, Verbeke, Vipond, Corfe, & Owen, 2015; Mukhopadhya, Hansen, El-Omar, & Hold, 2012).

Importantly, accumulating evidence indicates that microbiome structure reflects an individual’s ecological and functional readiness for life in the wild. Across mammals, captivity is associated with reproducible shifts toward simplified, unstable microbial communities that differ from wild-type configurations, often accompanied by the loss of key fiber-degrading and short-chain fatty acid–producing taxa (Jonathan B. Clayton et al., 2018; Frankel, Mallott, Hopper, Ross, & Amato, 2019; McKenzie et al., 2017). Such captivity-associated dysbiosis has been linked to reduced survival and compromised fitness following reintroduction, highlighting the microbiome as a sensitive indicator of latent physiological deterioration not detectable by conventional clinical screening (Bahrndorff, Alemu, Alemneh, & Lund Nielsen, 2016; Kenyon et al., 2014). Together, these findings position the microbiome as a missing dimension of release readiness, integrating metabolic, immunological, and ecological components of host health.

Slow lorises (*Nycticebus* and *Xanthonycticebus*: Nekaris & Nijman, 2022), including the Critically Endangered Javan slow loris (*N. javanicus*), provide a particularly relevant model for examining microbiome-mediated constraints on release readiness. These nocturnal primates are heavily targeted by the illegal pet trade, in which dental clipping is commonly performed to eliminate the risk of venom bites and facilitate handling. This procedure is typically performed without sterile technique, exposing the pulp cavity and frequently leading to infection, periodontitis, and eventual veterinary extraction upon confiscation (Guno et al., 2022; Moore et al., 2014; Nekaris & Starr, 2015). However, dental trauma represents only one component of the cumulative physiological burden experienced by rehabilitating individuals, which may also include prolonged captivity, altered diet, repeated medical intervention, and chronic health impairment.

Beyond the immediate injury, clipping-induced dental pathology can generate lasting ecological disturbance within the oral cavity. Loss of tooth-associated microhabitats and gingival restructuring alter local physicochemical conditions and biofilm architecture, potentially reshaping oral microbial communities (Dewhirst et al., 2010; Maki, Kazmi, Barb, & Ames, 2021; Mark, Rossetti, Rieken, Dewhirst, & Borisy, 2016). Impaired feeding mechanics and modified dietary regimes further influence substrate availability for both oral and gut microbes (Starr & Nekaris, 2013). These captivity-associated pressures are expected to propagate along the oral–gut axis (Amato et al., 2013; Jonathan B. Clayton et al., 2018; Flint et al., 2012), with downstream implications for metabolic efficiency and immune regulation (Mukhopadhya et al., 2012; Pickard et al., 2017). Importantly, such microbial restructuring may arise not solely from dental trauma, but from the combined effects of injury, rehabilitation duration, and broader medical history. In rehabilitating *N. javanicus*, behavioral traits such as activity budgets and boldness have recently been proposed to predict release readiness (Langgeng et al., 2025), establishing a behavioral framework for assessing release readiness. Yet individuals are typically classified as release candidates or non-candidates based on a combination of physical condition, behavior, and long-term prognosis, with severe dental trauma being a primary reason for permanent retention. Despite growing recognition of microbiome-mediated physiological resilience across mammals, no study has yet evaluated whether microbial communities capture complementary dimensions of alleged release readiness in this species.

Building on the view of the microbiome as an ecological organ of the host (Turnbaugh et al., 2007), evidence that captivity consistently erodes wild-type microbial configurations (Jonathan B. Clayton et al., 2018; Frankel et al., 2019; Hayakawa et al., 2018), and demonstrations that dysbiosis can compromise post-release survival (Kenyon et al., 2014), we posit that microbiome structure should reflect an individual’s latent ecological readiness for life in the wild. Because dental condition, medical burden, and duration of rehabilitation jointly influence diet, inflammation, and environmental exposure, microbiome composition may integrate these factors into a tractable biological signal. (Cabana et al., 2019; Maki et al., 2021).

Thus, in this study, we tested the hypothesis that microbiome structure constitutes an integrative ecological component of release readiness in rehabilitating *N. javanicus*. Specifically, we examined whether (i) oral and gut microbiomes form distinct, body site–specific ecological assemblages; (ii) microbiome composition differs between individuals classified as release candidates and non-candidates based on integrative rehabilitation criteria; and (iii) progression through soft-release is accompanied by partial restoration of microbial diversity and metabolic function toward wild-type configurations. We predicted that non-candidates would exhibit qualitative restructuring of microbial communities and functional profiles consistent with constrained metabolic or immunological flexibility, whereas release candidates would retain more stable and biosynthesis-oriented microbial configurations. Finally, we evaluated whether short-term exposure to semi-natural conditions during soft-release is accompanied by detectable microbial reorganization, or whether microbiome structure remains stable across this transition. By explicitly testing these predictions, our study evaluates whether microbiome-based metrics can reveal latent physiological variation aligned with rehabilitation and release decisions.

## METHODS

### Ethics Statement

This research was conducted in accordance with the Guidelines for the Care and Use of Non-human Primates and the Guidelines for Field Research on Non-human Primates established by the Center for the Evolutionary Origins of Human Behavior, Kyoto University. We obtained ethical approval and permissions from the Field Research Committee of the Wildlife Research Center, Kyoto University, as well as the Indonesian National Research and Innovation Agency (BRIN, Ref. no.: 009/KE.02/SK/01/2024). We also obtained the permission to collect biological samples from nationally protected species from the Director General of Natural Resources and Ecosystem Conservation (Dirjen KSDAE, Ref. no.: SK.127/KSDAE/SETKSDAE/KSA2/6/2024). All procedures adhered to the American Society of Primatologists (ASP) Principles for the Ethical Treatment of Non-Human Primates and the ASP Code of Best Practices for Field Primatology.

### Study site, subjects, and sample collection

We conducted sample collection from June to October 2024 at the Yayasan Inisiasi Alam Rehabilitasi Indonesia (YIARI). We selected 19 adult rehabilitating *N. javanicus*, (12 females and 7 males). The rehabilitation team classified ten individuals (7 females, 3 males) as release candidates (hereafter: candidates) based on assessment on oral and general health, medical history, behavioral performance, and overall rehabilitation trajectory culminating in release. The remaining nine individuals (5 females, 4 males) were designated non-releasable (hereafter: non-candidates) and retained permanently at the center because combinations of dental modification (tooth removal occurred in 4 of 10 candidates and all non-candidates), chronic or recurrent medical conditions, and/or physical impairment were judged to limit long-term survival in the wild. It is important to note that we determined release candidacy independent of this study. To clarify the structure underlying this classification, we conducted exploratory logistic regression analyses with candidacy as the response variable. Rehabilitation duration, dental condition, and medical history emerged as the strongest contributors (McFadden pseudo-R²: 0.31–0.84). Candidates had spent 5–32 months in rehabilitation (mean ± SD: 11.8 ± 8.9) and an additional two weeks in a soft-release enclosure before release into Mount Halimun-Salak National Park, West Java. In contrast, non-candidates had resided at the center for 29–145 months (84.4 ± 35.0). Exact chronological age and origins were unknown, as individuals were confiscated from trade or rescue situations without reliable background records.

In total, we collected 26 fecal samples, i.e., 10 from candidates during the pre-release phase, 7 from candidates during the soft-release phase, and 9 from non-candidates; as well as 18 saliva samples (9 each from candidate and non-candidate groups) (Table 1). We collected fecal samples opportunistically, less than 30 minutes upon defecation and obtained saliva samples during routine health check. We stored all samples in lysis buffer (0.5% SDS, 100 mM EDTA pH 8.0, 100 mM Tris-HCl pH 8.0, and 10 mM NaCl) at room temperature (Jose et al., 2024). We provided further details on individuals background (oral condition, medical history, year the individuals admitted to the rehabilitation center, and social housing conditions) and sampling history in Table S1.

**Table 1.** Putative pathogenic or opportunistic taxa detected in the microbiomes of rehabilitating *N. javanicus*. Listed taxa include genera, families, or species previously reported to be associated with oral or gut dysbiosis, inflammation, or opportunistic expansion under altered host or environmental conditions, rather than obligate pathogens.

| **Taxa** | **Enriched individuals** | **Sites** |
| --- | --- | --- |
| *Porphyromonas* | Candidates | Oral |
| *Fusobacterium nucleatum* | Non-candidates | Oral |
| **Pasteurellaceae** | Non-candidates | Oral |
| *Staphylococcus* | Non-candidates | Oral |
| *Morganella morganii* | Non-candidates | Oral |
| *Providencia* | Non-candidates | Oral |
| *Desulfovibrio* | Non-candidates | Oral and Gut |
| *Enterococcus* | Non-candidates | Gut |
| *Sutterella* | Non-candidates | Gut |
| *Fusobacterium varium* | Non-candidates | Gut |

### 16S rRNA gene sequencing

We extracted DNA from fecal samples using the QIAamp DNA Stool Mini Kit (QIAGEN GmbH, Hilden, Germany) following the manufacturer’s protocol with minor modifications describedby Lee et al. (2023). Library preparation and sequencing were performed by commercial company (PT. Genetika Science Indonesia). The V3 – V4 hypervariable regions of the bacterial 16S rRNA gene were targeted using the primer pair 341F (Forward: CCTAYGGGRBGCASCAG) and 806R (Reverse: GGACTANNGGGTATCTAAT).

Sequencing was performed on an Illumina MiSeq platform using paired-end sequencing We quality-filtered and denoised raw reads and inferred amplicon sequence variants (ASVs) using the DADA2 pipeline, then assigned taxonomy against the SILVA reference database (silva_nr99_v138.2).

### Data analyses

We calculated alpha and beta diversity using the R packages phyloseq (McMurdie & Holmes, 2013) and microbiome (Lahti & Shetty, 2017). We quantified alpha diversity using observed ASV richness and Shannon diversity. To test the effects of release candidacy and rehabilitation phase, we fitted generalized linear mixed models (GLMMs) in glmmTMB (Brooks et al., 2017), with richness or Shannon diversity as response variables, candidacy and phase as fixed effects, and sex as a covariate. We included individual identity as a random intercept to account for repeated sampling. Because rehabilitation phase was structurally associated with rehabilitation duration, we did not include duration in the same models. Instead, we assessed associations between rehabilitation duration and alpha diversity within candidate and non-candidate groups using Spearman rank correlations. We calculated weighted and unweighted UniFrac distances to assess beta diversity and tested group differences using PERMANOVA in vegan (Oksanen et al., 2025). We identified differentially abundant taxa using Linear Discriminant Analysis (LDA) effect size (LEfSe)(Segata et al., 2011), implemented in microbiomeMarker (Cao et al., 2022) applying a logarithmic LDA threshold of 2.0 (p < 0.05). We inferred functional profiles using Reconstruction of Unobserved States (PICRUSt2)(Douglas et al., 2020). We used the DHARMa package (Hartig, 2025) to test for homoscedasticity and overdispersion in the models. Finally, we ran ANOVA to evaluate statistical strength of all constructed models against null models in which all fixed effects terms had been removed.

## RESULTS

### Overall

After quality filtering, 2,389,127 high-quality reads were obtained from 44 samples (26 fecal and 18 saliva), with a mean of 54,298 ± SD 10,104 reads per sample. A total of 1,749 ASVs were detected across 15 phyla, 23 classes, 57 orders, 126 families, and 293 genera. Across body sites, 837 ASVs were unique to gut samples, 819 were unique to oral samples, and 93 were shared (Fig. 1A). Within gut microbiomes, candidate and non-candidate individuals harbored 327 and 349 unique ASVs, respectively, and shared 254 ASVs. Within oral microbiomes, 318 and 419 ASVs were unique to candidate and non-candidate individuals, respectively, with 175 ASVs shared between groups. Overall, the gut microbiome was dominated by Bacteroidota (58.4% ± 7.5%), Bacillota (24.8% ± 5.8%), and Actinomycetota (12.5% ± 4.0%) (Fig. 1B), whereas the oral microbiome showed a higher proportion of Pseudomonadota (60.3% ± 12.6%) and Bacillota (21.8% ± 7.0%), with Bacteroidota and Fusobacteriota present in lower abundances (8.0% ± 8.2% and 7.9% ± 8.4%, respectively) (Fig. 1C).

**Figure 1.**
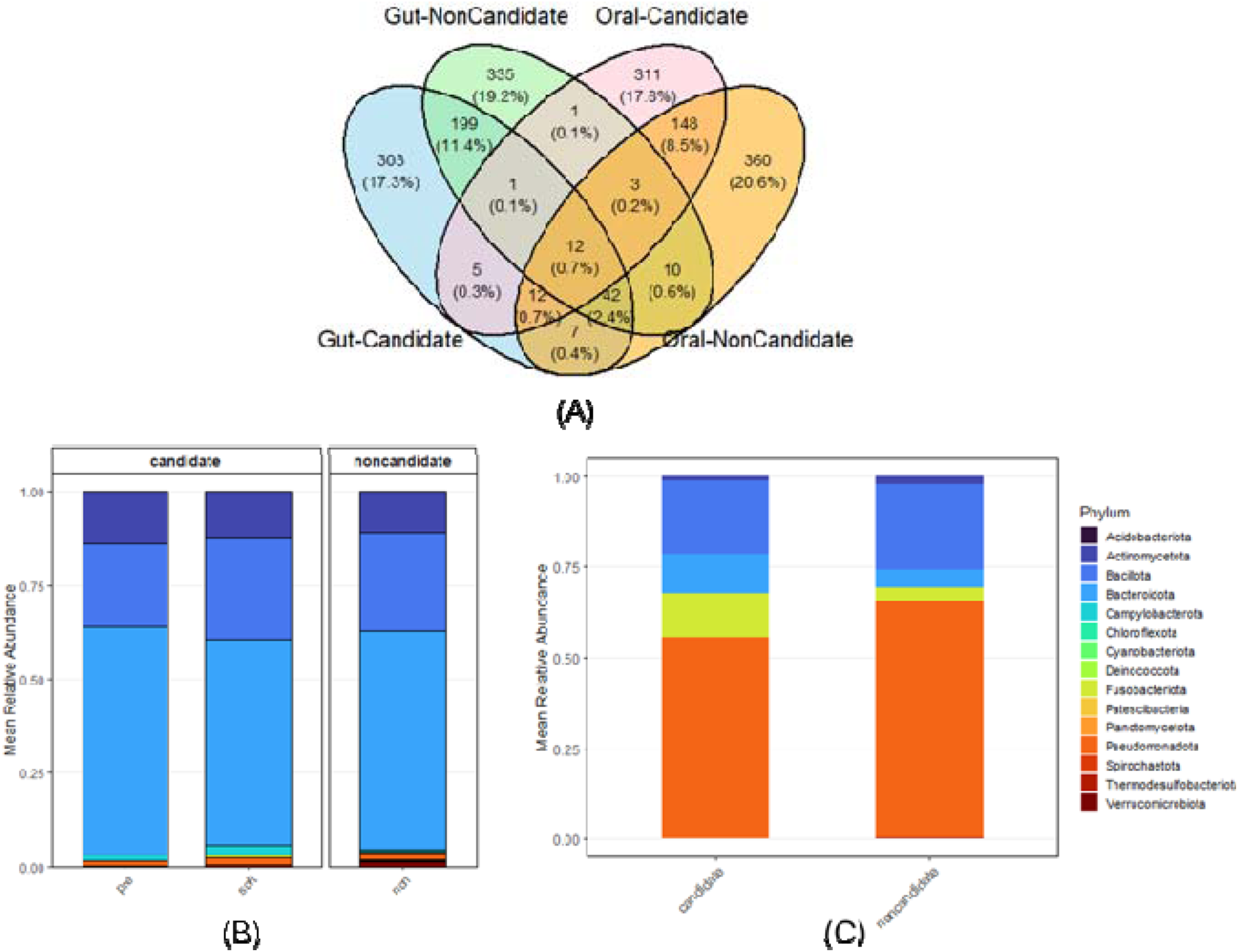
Taxonomic overview of gut and oral microbiomes in rehabilitating *N. javanicus*; (A) venn diagram showing the number of unique and shared amplicon sequence variants (ASVs) among gut–candidate, gut–non-candidate, oral–candidate, and oral–non-candidate groups; (B) mean relative abundance of bacterial phyla in gut microbiomes across candidate (pre-release and soft-release) and non-candidate individuals; (C) mean relative abundance of bacterial phyla in oral microbiomes of candidate and non-candidate individuals. Only phyla with a mean relative abundance ≥1% in at least one group are shown.

**Figure 2.**
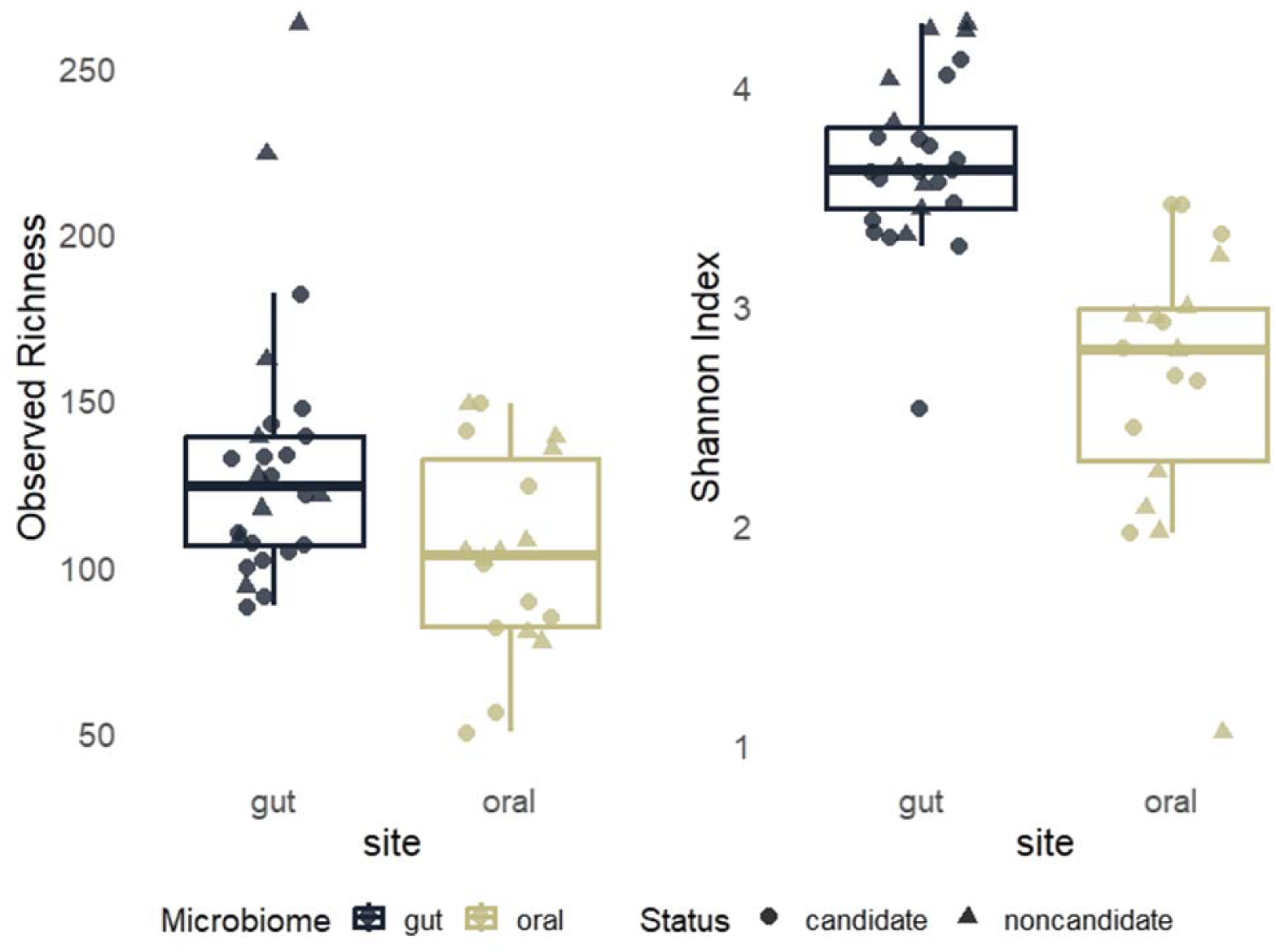
Alpha diversity of gut and oral microbiomes; observed ASV richness (left) and Shannon diversity index (right) for gut and oral microbiomes. Points represent individual samples; boxplots indicate median and interquartile range. Circles denote candidate individuals and triangles denote non-candidate individuals.

### Alpha diversity

#### Gut vs oral microbiomes

The gut microbiome exhibited significantly higher alpha diversity than the oral microbiome. Both observed richness (p = 0.004; Table S2) and Shannon diversity index (p < 0.001; Table S3) were significantly lower in oral samples. Neither sex nor rehabilitation duration influenced either diversity metric, indicating consistently greater taxonomic richness and evenness in gut compared to oral communities.

#### Gut microbiome

Within gut microbiomes, observed richness did not differ between candidate and non-candidate individuals (p = 0.27; Table S4), across rehabilitation duration (p >0.05; Table S5), or between pre- and soft-release phases (p = 0.93; Table S6). However, sex significantly affected observed richness in the pre-soft-release model. Shannon diversity showed only marginal differences between candidate and non-candidate individuals (p = 0.051; Table S7) and was not associated with rehabilitation duration (p > 0.05; Table S8) or release phase (p = 0.785; Table S9). Collectively, gut alpha diversity was largely stable across rehabilitation status, duration, and release phase, with persistent sex-related variation.

#### Oral microbiome

In the oral microbiome, neither observed richness nor Shannon diversity differ between candidate and non-candidate individuals (richness: p = 0.295; Table S10; Shannon: p = 0.135; Table S12), nor were they associated with rehabilitation duration (all p > 0.05; Table S11, S13). These results indicate stable oral alpha diversity across rehabilitation status, sex, and captivity duration.

### Beta Diversity

#### Gut vs oral microbiomes

Both unweighted and weighted UniFrac distances showed strong differentiation by body site (p = 0.001; Table S14-S15, Fig. 3). Microbiome composition was also influenced by rehabilitation status, whereas sex had no significant effect. These results indicate that body site and release status are primary determinants of microbial community structure.

**Figure 3.**
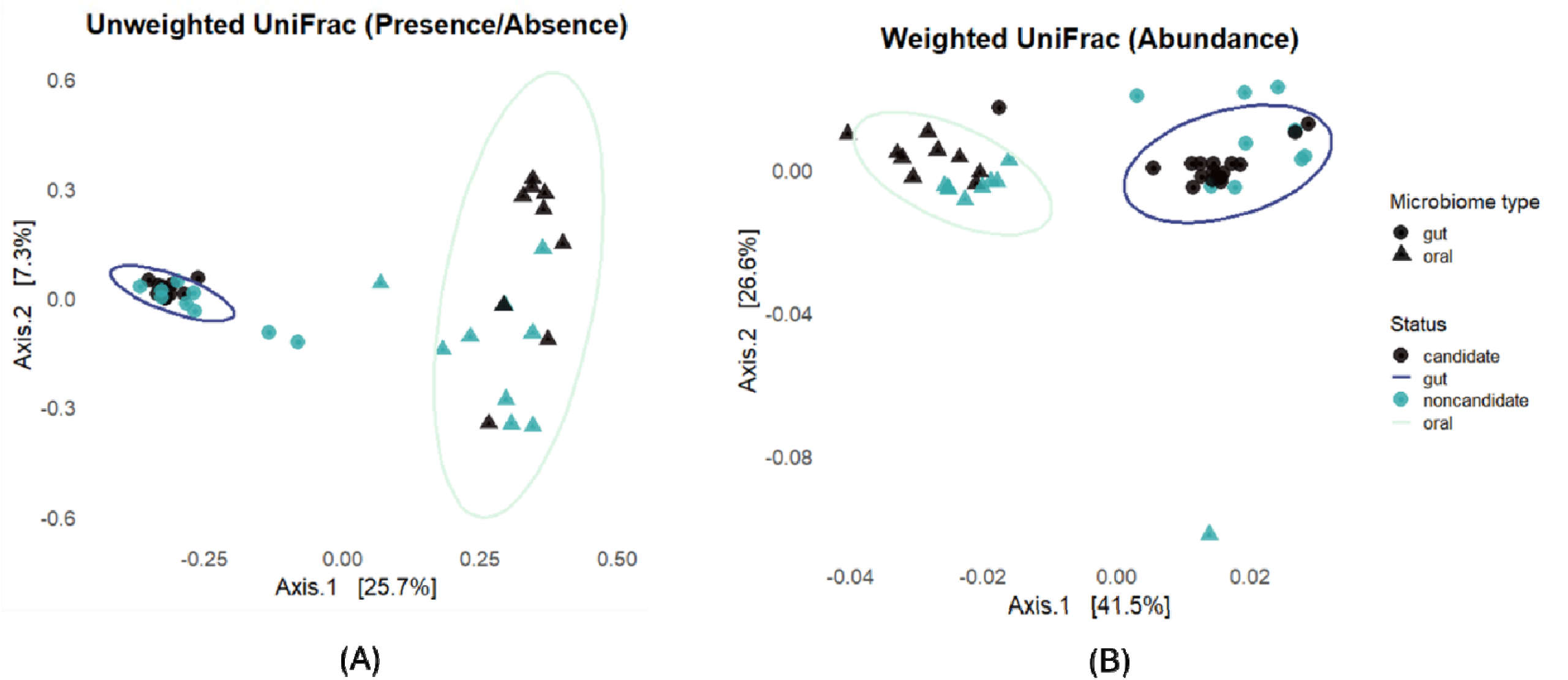
Beta diversity of gut and oral microbiomes; PCoA based on unweighted UniFrac (presence/absence; left) and weighted UniFrac (abundance-weighted; right) distances. Samples are colored by microbiome type (gut vs. oral) and shaped by release candidacy. Ellipses represent 95% dispersion of samples within each group.

**Figure 4.**
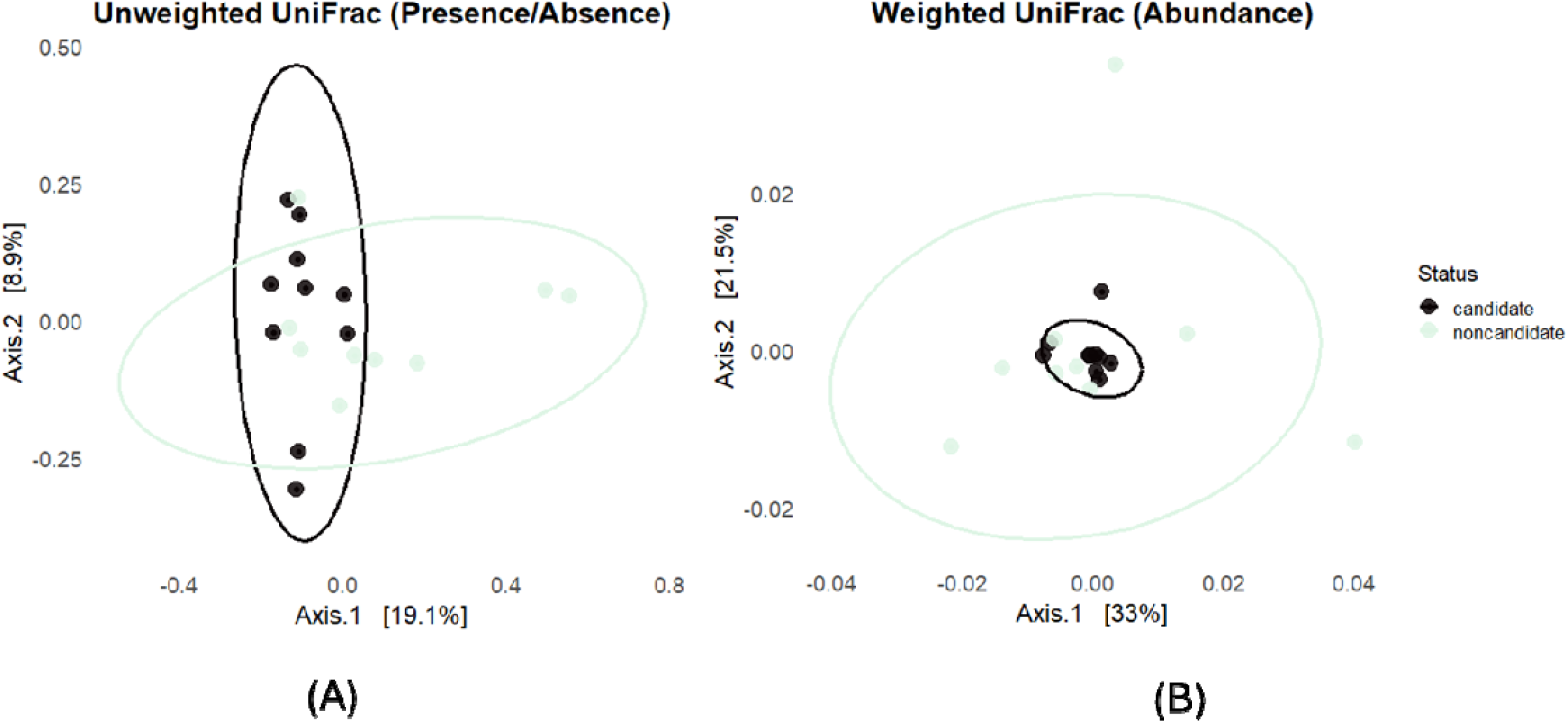
Gut microbiome composition by release candidacy; PCoA plots of gut microbiome communities based on unweighted UniFrac (left) and weighted UniFrac (right) distances, comparing candidate and non-candidate individuals. Ellipses represent 95% dispersion of samples within each group.

**Figure 5.**
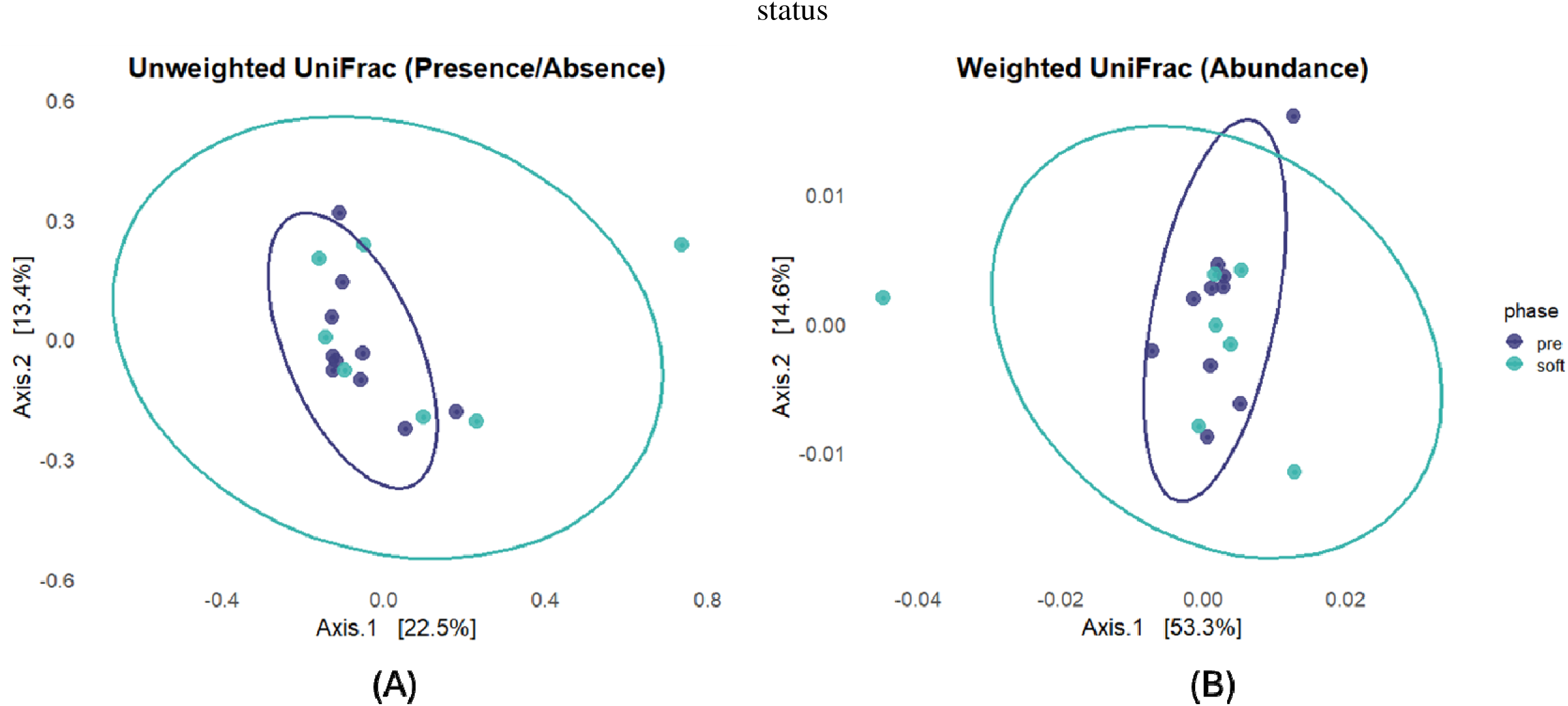
Gut microbiome composition across translocation phases; PCoA plots of gut microbiome communities based on unweighted UniFrac (left) and weighted UniFrac (right) distances, comparing pre-release and soft-release phases among candidate individuals. Ellipses represent 95% dispersion of samples within each group.

**Figure 6.**
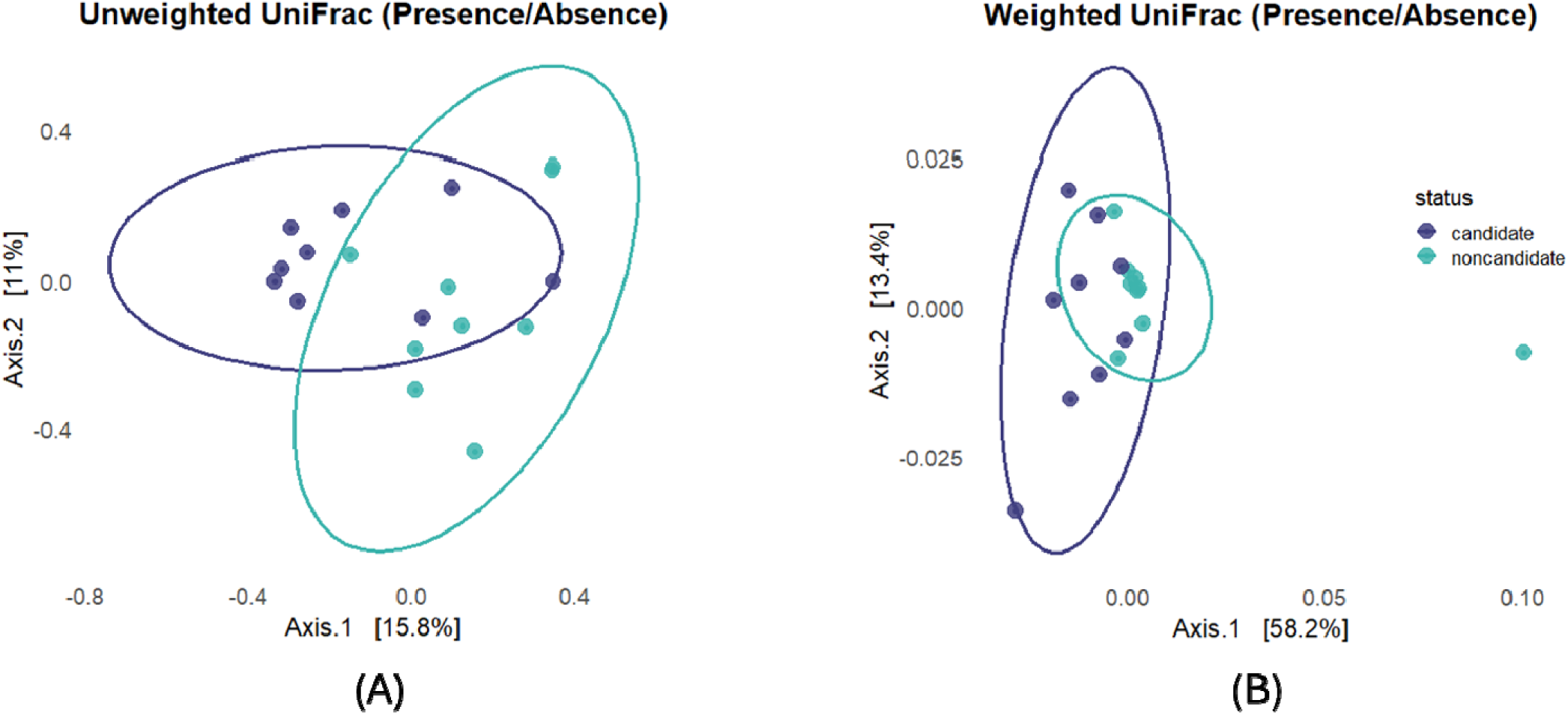
Oral microbiome composition by release candidacy; PCoA plots of oral microbiome communities based on unweighted UniFrac (right) and weighted UniFrac (left) distances, comparing candidate and non-candidate individuals. Ellipses represent 95% dispersion of samples within each group.

#### Gut microbiome

Unweighted UniFrac distances revealed a significant effect of candidate status (p = 0.032; Table S16), whereas weighted UniFrac distances did not (p = 0.996; Table S17), indicating that gut community differences between candidate and non-candidate individuals are driven primarily by presence/absence patterns rather than relative abundances. In contrast, neither unweighted nor weighted UniFrac distances differed between pre- and soft-release phases (unweighted: p = 0.57; weighted: p = 0.73; Table S18-S19), suggesting overall compositional stability across release phases.

#### Oral microbiome

Both unweighted and weighted UniFrac distances revealed significant differentiation between candidate and non-candidate individuals (all p < 0.01; Table S20-S21), indicating robust compositional divergence in oral microbial communities irrespective of abundance weighting.

### Enriched Microbial Taxa

#### Gut vs oral microbiomes

LEfSe analysis revealed clear taxonomic segregation between gut and oral microbiomes (LDA ≥ 2, p < 0.05; Figure S4). The gut microbiome was enriched in Bacteroidota, Actinomycetota, Spirochaetota, and Campylobacterota. In contrast, the oral microbiome was enriched in Pseudomonadota and Fusobacteriota (p < 0.001).

#### Gut microbiome: candidate vs non-candidate

Only a single genus, *Olsenella*, was enriched in candidate individuals (p = 0.039). In contrast, non-candidates exhibited enrichment of multiple taxa associated with opportunistic expansion and altered host–microbe interactions, including *Enterococcus* (p = 0.023), *Sutterella* (p = 0.034), *Sphaerochaeta* (p = 0.038), and the Rikenellaceae RC9 gut group (p = 0.008). At the species level, *Bacteroides uniformis* and *Fusobacterium varium* were enriched in non-candidates (both p = 0.022), together with additional genera including *Thomasclavelia,* UCG-004, *Lachnoclostridium*, and unclassified Tannerellaceae (Fig. 7A).

**Figure 7.**
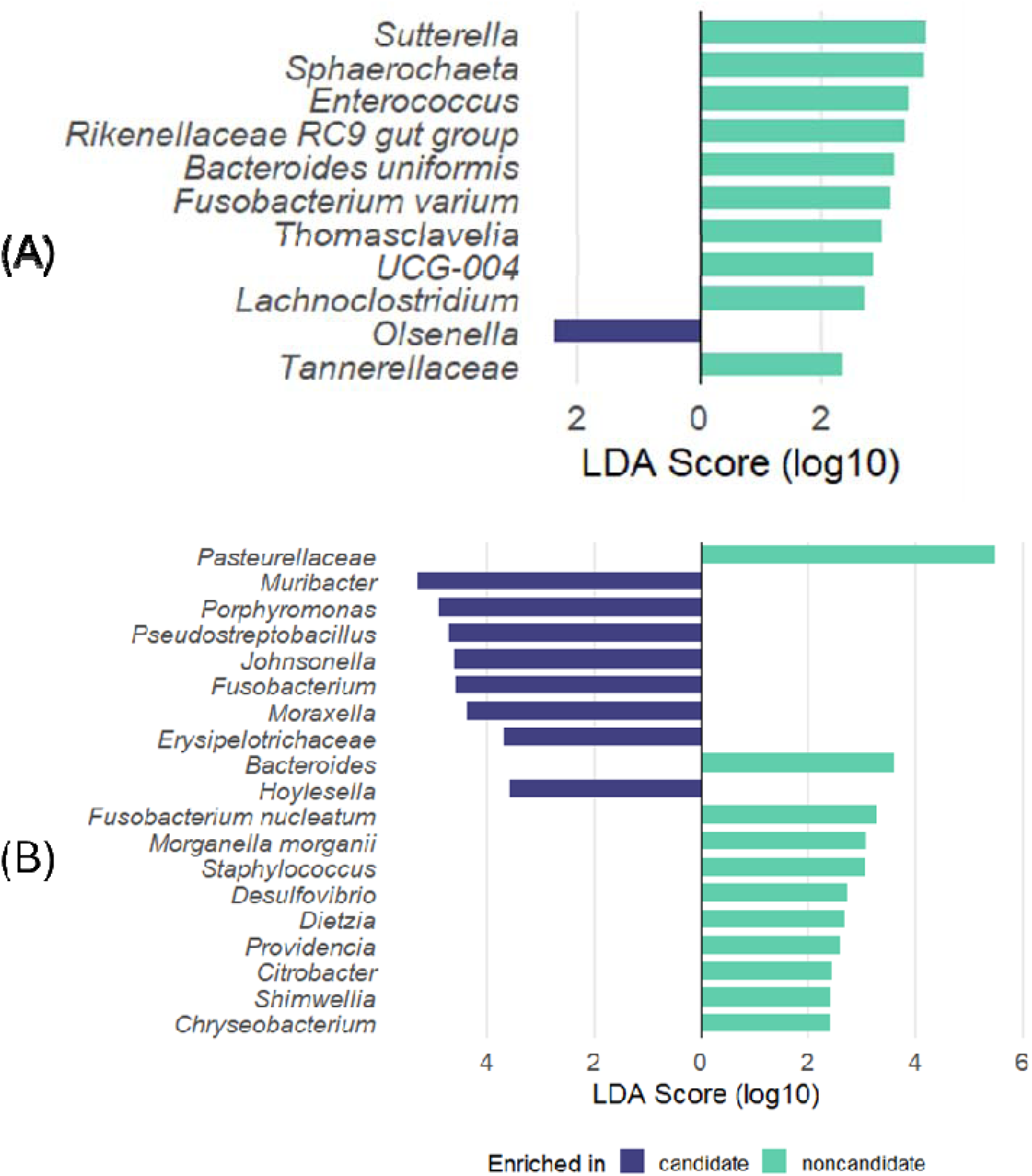
Differentially enriched taxa in gut and oral microbiomes; linear discriminant analysis effect size (LEfSe) results showing taxa significantly enriched between candidate and non-candidate individuals: (A) gut microbiome and (B) oral microbiome; bars represent log -transformed LDA scores; taxa enriched in candidates are shown on the left and those enriched in non-candidates on the right.

#### Release phase: pre- vs soft-release

Pre-release individuals were enriched in Prevotellaceae UCG-001 (p = 0.025), whereas soft-release individuals were enriched in *Lacticaseibacillus* (p = 0.049). No additional taxa differed significantly between phases.

#### Oral microbiome: candidate vs non-candidate

Candidate individuals were enriched in typical oral commensals and biofilm-associated taxa, with *Muribacter* showing the strongest enrichment, followed by *Porphyromonas*, *Pseudostreptobacillus*, *Johnsonella*, *Fusobacterium* (genus level), *Moraxella*, *Hoylesella*, and unclassified Erysipelotrichaceae (all p < 0.05). In contrast, non-candidates were characterized by enrichment of enteric and opportunistic taxa, with Pasteurellaceae showing the strongest discriminatory power (LDA = 5.46, p = 0.001), including *Citrobacter*, *Morganella morganii*, *Providencia*, *Shimwellia*, and *Fusobacterium nucleatum* (all p < 0.05; Fig. 7B).

#### Functional metagenomic profiles

LEfSe identified 16 differentially abundant functional pathways in the gut microbiome between groups (p < 0.05). Candidate individuals were enriched in multiple biosynthetic and energy metabolism pathways, including TCA cycle VII (acetate producers), TCA cycle VI (obligate autotrophs), the superpathway of thiamin diphosphate biosynthesis II, NAD salvage pathway II, L-lysine biosynthesis I, L-histidine degradation I, C4 photosynthetic carbon assimilation cycle, purine nucleobase degradation I (anaerobic), CMP-pseudaminate biosynthesis, and 2-methylcitrate cycle I (Fig. S5). In contrast, non-candidate gut microbiomes were enriched in lipid- and nitrogen-related pathways, including stearate biosynthesis II (bacteria and plants), gondoate biosynthesis (anaerobic), cis-vaccenate biosynthesis, putrescine biosynthesis IV, and nitrate reduction VI (assimilatory).

In the oral microbiome, candidate individuals were enriched in multiple biosynthetic and energy metabolism pathways, including starch biosynthesis, the reductive TCA cycle I, homoalactic fermentation, UMP biosynthesis, thiamin salvage IV, pyruvate fermentation to acetate and lactate II, L-isoleucine biosynthesis I, NAD salvage pathway II, heme biosynthesis, coenzyme A biosynthesis, and multiple glycolysis and aerobic respiration pathways (Fig. S6). In contrast, non-candidate oral microbiomes were enriched in degradation-associated pathways, including sugar alcohol degradation, cyclic polyol degradation, aromatic compound degradation, and multiple nucleoside and nucleotide degradation pathways, as well as lipid biosynthesis pathways (stearate biosynthesis II, phosphatidylglycerol biosynthesis I and II, gondoate biosynthesis) and additional pathways including L-histidine biosynthesis and the superpathway of menaquinol-8 biosynthesis I.

## DISCUSSION

This study characterizes oral and gut microbiomes in rehabilitating Javan slow lorises (*N. javanicus*) and shows that these compartments represent distinct physiological niches. Beyond this anatomical structuring, microbiome composition differed between release candidates and non-candidates, with the strongest divergence in the oral microbiome. Gut differences were more modest and primarily driven by presence–absence patterns. Short-term soft-release did not induce detectable microbial restructuring. Together, these findings suggest that microbiome profiles capture physiological variation aligned with rehabilitation status.

### Microbiome compartmentalization defines physiological niches

In this study, alpha diversity was consistently higher in the gut than in the oral cavity, as indicated by both observed richness and Shannon diversity. This pattern reflects the pronounced ecological differentiation between gastrointestinal and oral habitats reported across host-associated microbiome studies, although the magnitude and direction of diversity differences vary depending on host taxa, body site, and metrics used (Donaldson, Lee, & Mazmanian, 2016; Human Microbiome Project, 2012; Maki et al., 2021). Beta diversity analyses further demonstrated that body site was the strongest determinant of microbial community composition, exceeding the explanatory power of rehabilitation status or individual condition. This finding aligns with comparative work in primates and other mammals identifying anatomical compartmentalization as a dominant organizing principle of host-associated microbiomes (Asangba et al., 2022; Costello, Stagaman, Dethlefsen, Bohannan, & Relman, 2012; Lloyd-Price, Abu-Ali, & Huttenhower, 2016).

These patterns are consistent with fundamental physicochemical contrasts between compartments. The gut is a largely low-oxygen, nutrient-rich environment with prolonged transit times and extensive mucosal surface area, supporting dense communities dominated by obligate anaerobes specialized in fermentation and complex carbohydrate metabolism (Donaldson et al., 2016; Flint et al., 2012; Segata et al., 2011). In contrast, the oral cavity is a dynamic habitat shaped by fluctuating oxygen levels, mechanical disturbance during feeding and grooming, and continuous exposure to salivary antimicrobials. These factors favor biofilm-forming and facultatively anaerobic taxa organized along steep microscale gradients, resulting in community structures compositionally distinct from those of the gut gut (Avila, Ojcius, & Yilmaz, 2009; Dewhirst et al., 2010; Maki et al., 2021; Mark et al., 2016).

Although a limited subset of ASVs was shared between compartments, this overlap likely reflects transient taxa introduced through swallowing, generalists tolerating multiple mucosal environments, or contamination. Oral microbes are ingested in substantial numbers daily, and some may survive gastrointestinal transit, contributing to low-level cross-compartment similarity (Costello et al., 2012; Human Microbiome Project, 2012; Leao et al., 2023). Nonetheless, the predominance of site-specific ASVs underscores strong compartmental specialization and limited sustained exchange under typical physiological conditions (Donaldson et al., 2016; Lloyd-Price et al., 2016).

### Gut microbiome stability and qualitative restructuring across release candidacy

Contrary to the assumption that healthier microbiomes are necessarily more diverse, non-candidate *N. javanicus* exhibited marginally higher Shannon diversity in gut communities, while observed richness did not differ significantly between groups. Although this trend did not reach conventional significance thresholds and should be interpreted cautiously givenmodest sample size, increased diversity does not inherently indicate improved microbiome condition. Perturbed communities can exhibit comparable or even elevated diversity relative to more stable states, challenging simplistic “diversity equals health” paradigms (Carding et al., 2015; Zaneveld, McMinds, & Vega Thurber, 2017).

Consistent with this perspective, beta diversity analyses revealed modest but significant differentiation in unweighted UniFrac distances between candidates and non-candidates, whereas weighted UniFrac metrics showed no effect. This pattern indicates restructuring driven primarily by gains or losses of low-abundance taxa rather than shifts in dominant lineages (C. Lozupone, Lladser, Knights, Stombaugh, & Knight, 2011; Zaneveld et al., 2017). Such presence–absence divergence aligns with the “Anna Karenina principle,” whereby stressed microbiomes diverge idiosyncratically while stable communities remain comparatively similar (Zaneveld et al., 2017) .

A plausible mechanism underlying this pattern is reduced colonization resistance in disturbed gut ecosystems. In stable communities, dominant commensals suppress opportunistic colonizers through niche competition, antimicrobial production, and immune regulation (Kamada et al., 2013; Pickard et al., 2017). When these regulatory processes weaken, additional taxa may establish without enhancing functional stability, potentially inflating diversity estimates (Buffie & Pamer, 2013; Carding et al., 2015). Although causal inference is not possible, the convergence of marginal alpha diversity trends with presence/absence–driven beta diversity differences suggests that qualitative restructuring, rather than diversity per se, may better reflect gut microbiome condition in rehabilitating *N. javanicus*.

### Oral microbiomes as sensitive indicators of long-term dental perturbation

In contrast to the gut, oral microbiome alpha diversity did not differ between candidate and non-candidate individuals, indicating comparable levels of taxonomic richness and evenness across groups. Similar stability in oral alpha diversity has been reported in other primate studies, even when oral community composition differs markedly under captivity or pathological conditions (Maki et al., 2021; Ni et al., 2022). Despite this apparent stability in alpha diversity, beta diversity analyses revealed strong and consistent compositional differentiation between candidate and non-candidate oral microbiomes across both unweighted and weighted UniFrac metrics. The concordance between presence/absence–based and abundance-weighted distances indicates broad restructuring of oral microbial communities rather than changes restricted to rare taxa (C. Lozupone et al., 2011). Notably, the magnitude of compositional divergence in the oral microbiome exceeded that observed in the gut, suggesting that oral microbial communities are particularly sensitive to perturbations.

Permanent anatomical alterations following dental trauma and chronic inflammatory processes such as periodontitis likely contribute to this pattern. Changes in gingival architecture, exposure of new tissue surfaces, and sustained inflammatory signaling may therefore create stable ecological niches that promote long-term reorganization of oral microbial communities. (Avila et al., 2009; Mark et al., 2016). Once established, such structurally constrained oral environments may limit recovery toward pre-injury microbial configurations, resulting in persistent and compositionally distinct oral microbiomes in non-candidate individuals (Dewhirst et al., 2010; Maki et al., 2021).

Whether the oral or gut microbiome functions as the more sensitive health indicator may be taxon-specific. In Bengal slow lorises (*Nycticebus bengalensis*), Ni et al. (2022) reported that oral microbiome composition was more sensitive to captivity duration than the fecal microbiome and suggested its utility for monitoring during translocation. This differs from broader primate comparisons indicating greater gut microbiome sensitivity to captivity (Asangba et al., 2022; Clayton et al., 2016). Together, these findings suggest that in slow lorises, the oral microbiome may provide complementary and potentially informative insights, especially given the high prevalence of dental trauma in confiscated individuals (Ni et al., 2022).

### Microbial composition reflects functional specialization across body sites

Consistent with the strong compartmentalization observed in diversity analyses, gut and oral microbiomes of rehabilitating *N. javanicus* differed markedly in dominant taxonomic composition. The gut microbiome was primarily composed of Bacteroidota, Bacillota, and Actinomycetota, phyla commonly reported as core components of mammalian gut ecosystems (Amato et al., 2018; Jonathan B. Clayton et al., 2018; Flint et al., 2012; Ley et al., 2008). Spirochaetota and Campylobacterota were present at lower relative abundances and contributed to overall diversity. In contrast, oral microbiomes were dominated by Pseudomonadota and Fusobacteriota, taxa commonly associated with mucosal biofilms and facultatively anaerobic oral environments in mammals (Leao et al., 2023; Mark et al. 2016). While functional traits cannot be inferred directly from higher-level taxonomy, the dominance of these groups is broadly consistent with metabolic specialization typical of host-associated microbial ecosystems (Flint et al., 2012; Thomas, Hehemann, Rebuffet, Czjzek, & Michel, 2011). More detailed functional differences between candidate and non-candidate individuals are therefore better interpreted in light of the predicted pathway analyses presented below.

### Contextualizing *N. javanicus* microbiomes within primates and captivity

Such taxonomic structure is broadly consistent with microbiome patterns reported across other primates and mammals relying on plant-derived substrates for energy and aligns with the obligate exudativory and fiber-rich dietary ecology of slow lorises (Amato et al., 2018; Bo et al., 2010; Jonathan B. Clayton et al., 2018; Ni et al., 2022; Cabana et al., 2019; Starr & Nekaris, 2013).

Comparisons with wild slow lorises indicate that captivity and rehabilitation can modify gut microbiome structure. Wild individuals often exhibit greater inter-individual variability and, in some cases, higher relative abundances of Actinomycetota (Bo et al., 2010; Cabana et al., 2019), whereas rehabilitating individuals under controlled conditions tend to display more homogenized communities. Similar captivity-associated homogenization has been reported in other primates and mammals and may also reflect increased exposure to human-associated microbes through caretaker contact and shared infrastructure (Alberdi, Martin Bideguren, & Aizpurua, 2021; J. B. Clayton et al., 2016; McKenzie et al., 2017).

Data on slow loris oral microbiomes remain limited. To date, only one study has characterized the oral microbiome of a *Nycticebus* species, focusing on captive *N. bengalensis* (Ni et al., 2022). That study reported oral communities dominated by Pseudomonadota, Bacillota, and Fusobacteriota, with confinement duration associated with increased abundance of taxa linked to oral pathology. The similar taxonomic composition observed here suggests broadly comparable oral microbiome structure across *Nycticebus* species under captive conditions.

### Taxonomic and functional signatures of dysbiosis in non-candidate individuals

#### Gut microbiome

Integration of taxonomic and predicted functional profiles suggests that gut microbiome differences between candidate and non-candidate *N. javanicus* reflect qualitative community restructuring rather than simple loss of diversity. Candidate individuals showed enrichment of a single genus (*Olsenella sp.*), whereas non-candidates exhibited enrichment of multiple taxa alongside broader shifts in predicted metabolic pathways, consistent with increased ecological instability rather than targeted functional optimization. *Olsenella sp.*, enriched in candidates, is commonly reported as a fermentative Actinomycetota member of mammalian gut communities and has been associated with anaerobic metabolism and carbohydrate fermentation in herbivorous or fiber-rich dietary contexts (Cabana et al., 2019; Flint et al., 2012). Its selective enrichment in candidates, together with predicted pathways related to central energy metabolism and biosynthesis, is consistent with relatively stable gut ecosystems structured around fermentation-based energy extraction.

In contrast, non-candidate individuals showed enrichment of a heterogeneous assemblage of gut-associated taxa, including *Enterococcus sp.*, *Sutterella sp.*, *Sphaerochaeta sp.*, members of the Rikenellaceae RC9 gut group, *Bacteroides uniformis*, and *Fusobacterium varium*. Several of these taxa have been reported to expand under perturbed or clinically altered gut conditions and have been associated with altered host–microbe interactions in mammals (Carding et al., 2015; Mukhopadhya et al., 2012). Others, such as Rikenellaceae RC9 and *B. uniformis*, are common commensals whose abundance is strongly shaped by diet and environmental context rather than pathogenicity *per se* (Cabana et al., 2019; Jonathan B. Clayton et al., 2018). These taxonomic shifts were accompanied by enrichment of predicted functional pathways related to polyamine metabolism, fatty acid biosynthesis, and nitrate reduction. Polyamines such as putrescine play roles in epithelial maintenance, but altered regulation of polyamine metabolism has also been linked to epithelial stress and inflammatory responses under disturbed gut conditions (Carding et al., 2015). Similarly, enrichment of nitrate respiration pathways is a well-documented microbial response to gut environments experiencing increased oxygen or nitrate availability, conditions often associated with inflammation or dysbiosis (Mukhopadhya et al., 2012).

Together, these coordinated taxonomic and functional patterns are consistent with gut microbial communities structured to tolerate disturbed or fluctuating conditions, rather than optimized for strict anaerobic fermentation. Such restructuring likely reflects cumulative pressures common in long-term rehabilitation, including dental impairment and associated dietary modifications, general health and repeated medical intervention, and prolonged captivity (Cabana et al., 2019; Ni et al., 2022).

#### Oral microbiome

Differences between candidate and non-candidate individuals were more pronounced in the oral microbiome than in the gut, with strong compositional differentiation accompanied by shifts in predicted functional profiles. Candidate individuals were enriched in several taxa commonly detected in mammalian oral biofilms, including *Muribacter sp.*, *Porphyromonas sp.*, *Pseudostreptobacillus sp.*, *Johnsonella sp.*, *Moraxella sp.*, and genus-level *Fusobacterium spp.*. Many of these taxa occur in both healthy and diseased oral communities, and their ecological roles are strongly context-dependent rather than intrinsically pathogenic (Dewhirst et al., 2010; Francavilla et al., 2014; Han, 2015).

In contrast, non-candidate oral microbiomes were characterized by enrichment of taxa more consistently associated with perturbed oral environments. Members of the family Pasteurellaceae showed the strongest discriminatory signal, a group that in humans and other mammals has been linked to oral and respiratory infections under dysbiotic or immunocompromised conditions (Batı, Sezer, Akyüz, & Kırmızıgül, 2024; Dewhirst et al., 2010; Human Microbiome Project, 2012; Mach, Baranowski, Nouvel, & Citti, 2021). Enrichment of *Fusobacterium nucleatum*, rather than genus-level *Fusobacterium*, is particularly notable given its established association with inflammatory and periodontal states across host species (Han, 2015). Non-candidates also exhibited enrichment of taxa typically classified as enteric bacteria (*Citrobacter sp.*, Morganella sp., *Providencia sp.*, *Shimwellia sp.*). The presence of enteric-associated taxa in the oral cavity has been linked to compromised mucosal barriers, altered compartmentalization, or increased microbial translocation in primates and other mammals (Costello et al., 2012; Human Microbiome Project, 2012). In addition, although extremely rare, coprophagy has been observed in *N. javanicus* at the study center (YIARI, unpublished data), and may represent a plausible behavioral route for the transient introduction of enteric taxa into the oral cavity. Alternatively, incidental introduction via diet, environmental exposure, or handling during feeding and sampling cannot be excluded.

Functionally, candidate oral microbiomes were enriched in predicted biosynthetic and central metabolic pathways, including nucleotide and amino acid biosynthesis, vitamin and cofactor salvage, and core energy metabolism. Such profiles are characteristic of metabolically active oral communities capable of sustained growth under fluctuating conditions (Maki et al., 2021). In contrast, non-candidate oral microbiomes showed enrichment of degradation- and recycling-oriented pathways, including those related to sugar alcohols, aromatic compounds, nucleosides, and lipids. Similar functional shifts have been reported in oral microbial communities experiencing environmental stress or altered substrate availability (Ni et al., 2022).

Taken together, these results indicate a functional shift from biosynthesis-oriented metabolism in candidates toward degradation- and persistence-oriented strategies in non-candidates, consistent with long-term restructuring of the oral microbiome following dental trauma.

### Gut microbiome stability across release phases

Despite exposure to novel environments during the transition from pre-release to soft-release, gut microbiome structure remained stable. Neither alpha diversity nor beta diversity differed significantly between phases, indicating limited short-term microbial restructuring. Such stability is consistent with previous observations that gut microbiomes can exhibit substantial resilience to moderate environmental or dietary perturbations, particularly when hosts retain functionally redundant core communities (Cabana et al., 2019; C. A. Lozupone, Stombaugh, Gordon, Jansson, & Knight, 2012).

Notably, the duration of soft-release exposure in this study (approximately two weeks) was relatively short, likely insufficient to induce large-scale compositional change. Previous work in primates has shown that pronounced gut microbiome shifts typically require sustained dietary or environmental alteration over longer timescales (Amato et al., 2013; Cabana et al., 2019). For example, Ni et al. (2021) demonstrated that an 8-week dietary transition in rehabilitating Bengal slow lorises (*N. bengalensis*) resulted in significant restructuring of gut microbial communities. In contrast, rehabilitation and soft-release diets in the present study were largely standardized; although individuals housed in forest enclosures may have had limited opportunities to forage insects opportunistically, core dietary provisioning remained similar. Thus, contrary to our initial prediction, short-term progression through soft-release was not accompanied by detectable restoration of gut microbial diversity or function, suggesting that microbiome recovery may require longer exposure to natural diets and environments or may be constrained by earlier captivity-associated perturbations.

### Outlook

#### Study limitations

Several limitations should be considered when interpreting these findings. First, sample sizes were modest (gut microbiome: 10 candidates and 9 non-candidates; oral microbiome: 9 individuals per group), limiting statistical power to detect subtle effects and constraining evaluation of interactions among sex, rehabilitation duration, and dietary variation. In addition, the largely cross-sectional design, with most individuals sampled at a single time point, precludes inference about causal relationships or temporal microbiome dynamics. Importantly, the classification of individuals as release candidates or non-candidates reflects an integrative rehabilitation assessment rather than a single biological variable. These categories therefore incorporate multiple correlated factors, including dental condition, medical history, and rehabilitation duration, that could not be disentangled with the present sample size. In particular, individuals in the two groups differed substantially in time spent in rehabilitation, making it difficult to isolate whether observed microbiome differences reflect candidacy status per se, cumulative captivity effects, or underlying health conditions. Longitudinal sampling across rehabilitation, soft-release, and post-release phases would provide stronger insight into microbiome trajectories and their association with health and survival.

Second, baseline microbiome data from wild *N. javanicus* or from individuals at the time of confiscation were unavailable. Consequently, it remains unclear whether microbiomes of release candidates reflect retention of wild-type communities or convergence toward a distinct, rehabilitation-associated state, and whether dysbiosis observed in non-candidates developed during captivity or represents pre-existing impacts of wildlife trade. Individuals in this study exhibited heterogeneous medical histories, including dental pathology, recurrent infections, gastrointestinal disturbances, malnutrition, and systemic conditions, all of which may plausibly influence microbial community structure through inflammation, altered diet, or medical intervention. Moreover, most mechanistic links between dysbiosis and pathology derive from humans or other mammals rather than lorisidae. We therefore interpret dysbiosis-like signatures as ecological indicators of altered host–microbe dynamics rather than evidence of diagnosed enteric disease. Establishing wild reference baselines, sampling at intake, and integrating microbiome profiles with structured health assessments would be essential to clarify these relationships (Cabana et al., 2019; Ni et al., 2022).

Finally, host physiological data were not collected alongside microbiome profiles. Integrating microbiome analyses with immunological, inflammatory, or metabolic markers would strengthen mechanistic inference linking microbial structure to host condition. Previous work in humans and wildlife demonstrates that such integrative approaches substantially improve interpretation of microbiome variation in relation to health and disease risk (Bahrndorff et al., 2016; Lloyd-Price et al., 2016).

#### Implications for rehabilitation and conservation

Our findings add a complementary physiological dimension to existing behavioral frameworks used to assess release readiness in rehabilitating *N. javanicus*. Recent work suggested that activity budgets and boldness profiles provide valuable indicators of behavioral welfare and exploratory capacity relevant to post-release performance (Langgeng et al., 2025). Such metrics are practical, non-invasive, and well suited to rehabilitation settings, and they capture key aspects of how individuals interact with their environment.

The present study suggests that oral microbiome profiles may capture information not fully reflected in overt behavior. Differences between candidates and non-candidates cannot be attributed to dental trauma alone, as dental pathology, medical history, and rehabilitation duration are structurally intertwined. Although rehabilitation duration was not significantly associated with oral diversity or composition in our models, limited sample size and collinearity constrain causal interpretation, and additional factors such as prior antibiotic exposure or housing conditions may contribute. These microbial patterns may therefore represent a complementary layer of information, reflecting cumulative physiological history or altered recovery trajectories that are not readily detected through behavioral observation alone.

Taken together, integrating behavioral indicators (e.g. activity budgets and boldness) with microbiome-informed perspectives may improve the resolution of pre-release assessments. Behavioral metrics may reflect short-term adaptability and exploratory potential, whereas microbiome structure may capture longer-term physiological consequences of captivity, trauma, or dietary modification. However, given the modest sample size and cross-sectional design, these findings should be considered preliminary.

Additional longitudinal studies integrating microbiome profiles with standardized health and post-release survival data will be necessary to evaluate whether microbiome-informed metrics meaningfully improve release assessment in slow loris conservation (Langgeng et al., 2025).

## Acknowledgements

We would like to thank Kyoto University Wildlife Research Center (WRC), the Center for International Collaboration and Advanced Studies in Primatology (CICASP), Research Units for Exploring Future Horizons (Coevolution and Coexistence), and the Joint Research Program of WRC for facilitating our research, Yayasan Inisiasi Alam Rehabilitasi Indonesia (YIARI) Bogor board of directors and staff (especially Mastur, Ajo, Aconk, Ganyong, Pak Aki, Kojek) for their permission and assistance in the field, FMIPA-Biologi IPB University dean and staff. We also thanked the Indonesian National Research and Innovation Agency (BRIN), the Ministry of Environment and Forestry (KLHK), the Directorate General of Biodiversity Conservation (Dirjen KSDAE), and the West Java Natural Resources Conservation Center (BBKSDAE Jawa Barat) for the administrative assistance and permission to conduct research in Indonesia.

## Author contributions

AL: Conceptualization, methodology, investigation, formal analysis, writing – original draft, funding acquisition, project administration. WP: writing – review & editing, project administration, investigation. NPP: writing – review & editing, project administration, investigation. PR: writing – review & editing, project administration. KLS: writing – review & editing, project administration. RM: writing – review & editing, project administration. MS: conceptualization, writing – review & editing, supervision, funding acquisition. WL: formal analysis, writing – review & editing. AJJM: conceptualization, formal analysis, writing – review & editing, supervision, funding acquisition. IM: conceptualization, formal analysis, writing – original draft, supervision, funding acquisition

## Data availability

All data needed to evaluate the conclusions in the paper are present in the paper. Additional data related to this paper may be requested from the authors.

## Declarations

*Generative AI and AI-assisted technologies in the writing process*: During the preparation of this manuscript, the authors used ChatGPT-5.2 to refine the clarity and logical flow of the text. The authors carefully reviewed, corrected, and approved all content generated, and take full responsibility for the final published version.

*Conflict of interest*: The authors declare no competing interests.

