## Supplementary material for "Oral and gut microbiomes reveal potential physiological constraints on release readiness in rehabilitating Javan slow lorises": AL_Microbiome_supplementary_AJP.docx

Table S1. Study subject information and biological sample availability. Sex, release status (candidate or non-candidate), year of admission to the rehabilitation center (YIARI), and availability of biological samples collected from each individual are shown. “X” indicates that a sample was collected for the corresponding individual and sampling phase.

| ID | Sex | Status |  |  | Year admitted to YIARI |  | Biological samples collected | | |
| --- | --- | --- | --- | --- | --- | --- | --- | --- | --- |
|  |  |  | Oral condition | Past medical history |  | Housing condition | Pre-release  (fecal) | Soft-release (fecal) | Oral (saliva) |
| JS01 | Female | candidate | mild | yes | 2022 | solitary | X | X | X |
| JS02 | Female | candidate | good | yes | 2024 | group | X |  | X |
| JS03 | Female | candidate | mild | yes | 2023 | solitary | X | X | X |
| JS04 | Male | candidate | good | no | 2024 | solitary | X | X | X |
| JS05 | Female | candidate | good | no | 2022 | group | X |  | X |
| JS06 | Female | candidate | good | no | 2024 | group | X | X | X |
| JS07 | Female | candidate | mild | yes | 2023 | solitary | X | X | X |
| JS08 | Female | candidate | mild | no | 2024 | group | X | X |  |
| JS09 | Male | candidate | mild | no | 2024 | solitary | X | X | X |
| JS10 | Male | candidate | good | no | 2024 | group | X |  | X |
| JS11 | Female | non-candidate | bad | yes | 2016 | solitary | X |  | X |
| JS12 | Female | non-candidate | mild | yes | 2012 | solitary | X |  | X |
| JS13 | Male | non-candidate | bad | yes | 2017 | solitary | X |  | X |
| JS14 | Female | non-candidate | bad | yes | 2022 | solitary | X |  | X |
| JS15 | Male | non-candidate | bad | yes | 2015 | solitary | X |  | X |
| JS16 | Female | non-candidate | bad | yes | 2015 | solitary | X |  | X |
| JS17 | Male | non-candidate | bad | yes | 2022 | solitary | X |  | X |
| JS18 | Male | non-candidate | bad | yes | 2018 | solitary | X |  | X |
| JS19 | Female | non-candidate | bad | yes | 2018 | solitary | X |  | X |

Dental condition: bad (multiple clipped/extracted teeth and/or chronic oral pathology); mild (limited clipping/extraction); good (dentition intact with no oral pathology).

Past medical history includes chronic or recurrent illness, infection (e.g. abscess, inflammation), injuries or trauma, metabolic or systemic disorders.

Table S2. Effects of body sites (gut and oral) on observed ASV richness. Estimates of negative binomial generalized linear mixed model (coefficients ± 95% CI) testing the influence of body site, sex and rehabilitation duration on observed ASV richness. Individual identity was included as a random effect.

Family: nbinom2 ( log )

Formula: Observed ~ sex + microbiome + monthsofrehab + (1 | id)

```
Conditional model:
                 Estimate Std. Error z value Pr(>|z|)
(Intercept)     4.8349930  0.0889056   54.38   <2e-16 ***
Female vs male  0.0862510  0.0950157    0.91   0.3640
Gut vs oral    -0.2630788  0.0923860   -2.85   0.0044 **
Rehab duration  0.0007799  0.0010678    0.73   0.4651
---
Signif. codes:  0 ‘***’ 0.001 ‘**’ 0.01 ‘*’ 0.05 ‘.’ 0.1 ‘ ’ 1
```

Table S3. Effects of body site on Shannon diversity. Results of the GLMM testing differences in Shannon diversity between gut and oral microbiomes, with sex included as a fixed effect and individual identity as a random effect.

Family: gaussian ( identity )

Formula: Shannon ~ sex + microbiome + monthsofrehab + (1 | id)

```
Conditional model:
                Estimate Std. Error z value Pr(>|z|)
(Intercept)     3.712619   0.147011  25.254  < 2e-16 ***
Female vs male  0.143175   0.159353   0.898    0.369
Gut vs oral     -1.027807   0.154837  -6.638 3.18e-11 ***
Rehab duration  -0.001399   0.001765  -0.793    0.428
```

Table S4. Effects of release candidacy on gut observed ASV richness. GLMM results testing the effect of release candidacy (candidate vs non-candidate) on observed ASV richness in gut microbiomes, with sex as a fixed effect and individual identity as a random effect.

Family: nbinom2 ( log )

Formula: Observed ~ sex + status + (1 | id)

Data: gut_alpha[gut_alpha$phase != "soft", ]

```
Conditional model:
                   Estimate Std. Error z value Pr(>|z|)
(Intercept)         4.74503    0.08612   55.10   <2e-16 ***
Female vs male      0.13117    0.11836    1.11    0.268
Candidate vs non-c  0.16369    0.11458    1.43    0.153
```

Table S5. Relationship between rehabilitation duration and gut observed ASV richness. Spearman correlation results assessing the association between rehabilitation duration and observed ASV richness in gut microbiomes.

```
S = 237.46, p-value = 0.2042
alternative hypothesis: true rho is not equal to 0
sample estimates:
       rho
-0.4391326

S = 184, p-value = 0.1475
alternative hypothesis: true rho is not equal to 0
sample estimates:
       rho
-0.5333333
```

Table S6. Effects of release phase on gut observed ASV richness. GLMM results testing differences in gut observed ASV richness between pre-release and soft-release phases, with sex as a fixed effect and individual identity as a random effect.

Family: gaussian ( identity )

Formula: Observed ~ sex + phase + (1 | id)

```
Conditional model:
           	 Estimate Std. Error z value Pr(>|z|)
(Intercept) 	113.6356     7.2235  15.731   <2e-16 ***
Female vs male 25.2146    10.8546   2.323   0.0202 *
Pre- vs soft-  0.8745    10.0494   0.087   0.9307
```

Table S7. Effects of release candidacy on gut Shannon diversity. GLMM results testing the effect of release candidacy on Shannon diversity in gut microbiomes.

Family: gaussian ( identity )

Formula: Shannon ~ sex + status + (1 | id)

```
Conditional model:
                   Estimate Std. Error z value Pr(>|z|)
(Intercept)         3.53094    0.09879   35.74   <2e-16 ***
Female vs male              0.12994    0.13671    0.95   0.3419
Candidate vs non-c  0.25732    0.13208    1.95   0.0514 .
```

Table S8. Relationship between rehabilitation duration and gut Shannon diversity. Spearman correlation results assessing the association between rehabilitation duration and gut Shannon diversity.

```
S = 103.93, p-value = 0.2924
alternative hypothesis: true rho is not equal to 0
sample estimates:
      rho
0.3701261

S = 176, p-value = 0.2125
alternative hypothesis: true rho is not equal to 0
sample estimates:
       rho
-0.4666667
```

Table S9. Effects of release phase on gut Shannon diversity. GLMM results testing differences in gut Shannon diversity between pre-release and soft-release phases.

Family: Gamma ( log )

Formula: Shannon ~ sex + phase + (1 | id)

```
Conditional model:
            	Estimate Std. Error z value Pr(>|z|)
(Intercept)  	1.25888    0.03433   36.67   <2e-16 ***
Female vs male 0.04734    0.05216    0.91    0.364
Pre- vs soft-  -0.01317    0.04829   -0.27    0.785
```

Table S10. Effects of release candidacy on oral observed ASV richness. GLMM results testing differences in observed ASV richness between candidate and non-candidate oral microbiomes.

Family: gaussian ( identity )

Formula: Observed ~ sex + status + (1 | id)

```
Conditional model:
                   Estimate Std. Error z value Pr(>|z|)
(Intercept)          97.632     10.612   9.200   <2e-16 ***
Female vs male       -1.895     13.932  -0.136    0.892
Candidate vs non-c   14.211     13.583   1.046    0.295
```

Table S11. Relationship between rehabilitation duration and oral observed ASV richness. Spearman correlation results assessing the association between rehabilitation duration and observed ASV richness in oral microbiomes.

```
S = 152.38, p-value = 0.4826
alternative hypothesis: true rho is not equal to 0
sample estimates:
       rho
-0.2698204

S = 142, p-value = 0.6436
alternative hypothesis: true rho is not equal to 0
sample estimates:
       rho
-0.1833333
```

Table S12. Effects of release candidacy on oral Shannon diversity. GLMM results testing differences in Shannon diversity between candidate and non-candidate oral microbiomes.

Family: gaussian ( identity )

Formula: Shannon ~ sex + status + (1 | id)

Data: oral_alpha

```
Conditional model:
                   Estimate Std. Error z value Pr(>|z|)
(Intercept)          2.8076     0.2095  13.401   <2e-16 ***
Female vs male       0.1596     0.2750   0.580    0.562
Candidate vs non-c  -0.4009     0.2682  -1.495    0.135
```

Table S13. Relationship between rehabilitation duration and oral Shannon diversity. Spearman correlation results assessing the association between rehabilitation duration and Shannon diversity in oral microbiomes.

```
S = 153.42, p-value = 0.468
alternative hypothesis: true rho is not equal to 0
sample estimates:
       rho
-0.2785242

S = 126, p-value = 0.9116
alternative hypothesis: true rho is not equal to 0
sample estimates:
  rho
```

-0.05

Table S14. PERMANOVA results for unweighted UniFrac distances across body sites. PERMANOVA results testing the effects of body site, release candidacy, and sex on microbiome composition based on unweighted UniFrac distances.

```
adonis2(formula = dist_uw ~ microbiome + status + sex, data = meta_nosoft, by = "margin")
           Df SumOfSqs      R2       F Pr(>F)
microbiome  1   3.3192 0.24447 11.5050  0.001 ***
status      1   0.4589 0.03380  1.5907  0.044 *
sex         1   0.2769 0.02039  0.9598  0.387
Residual   33   9.5206 0.70121
Total      36  13.5773 1.00000
```

Table S15. PERMANOVA results for weighted UniFrac distances across body sites. PERMANOVA results testing the effects of body site, release candidacy, and sex on microbiome composition based on weighted UniFrac distances.

```
adonis2(formula = dist_wu ~ microbiome + status + sex, data = meta_nosoft, by = "margin")
           Df SumOfSqs      R2       F Pr(>F)
microbiome  1 0.015578 0.36301 20.4777  0.001 ***
status      1 0.001507 0.03511  1.9805  0.059 .
sex         1 0.000845 0.01968  1.1101  0.356
Residual   33 0.025104 0.58499
Total      36 0.042914 1.00000
```

Table S16. PERMANOVA results for gut microbiomes using unweighted UniFrac distances. PERMANOVA results testing the effect of release candidacy on gut microbiome composition using unweighted UniFrac distances.

```
adonis2(formula = dist_uw_gut ~ status + sex, data = meta_gut, by = "margin")
         Df SumOfSqs      R2      F Pr(>F)
status    1   0.3147 0.07850 1.4443  0.032 *
sex       1   0.1912 0.04770 0.8776  0.719
Residual 16   3.4858 0.86958
Total    18   4.0086 1.00000
```

Table S17. PERMANOVA results for gut microbiomes using weighted UniFrac distances. PERMANOVA results testing the effect of release candidacy on gut microbiome composition using weighted UniFrac distances.

```
adonis2(formula = dist_wu_gut ~ status + sex, data = meta_gut, by = "margin")
         Df  SumOfSqs      R2      F Pr(>F)
status    1 0.0000282 0.00349 0.0611  0.996
sex       1 0.0006375 0.07883 1.3784  0.184
Residual 16 0.0073994 0.91505
Total    18 0.0080863 1.00000
```

Table S18. PERMANOVA results for gut microbiomes across release phases using unweighted UniFrac distances. PERMANOVA results testing differences in gut microbiome composition between pre-release and soft-release phases based on unweighted UniFrac distances.

```
adonis2(formula = dist_uw_gut2 ~ phase + sex, data = meta_gut2, by = "margin")
         Df SumOfSqs      R2      F Pr(>F)
phase     1   0.2042 0.05845 0.9333  0.572
sex       1   0.2265 0.06485 1.0354  0.407
Residual 14   3.0631 0.87686
Total    16   3.4933 1.00000
```

Table S19. PERMANOVA results for gut microbiomes across release phases using weighted UniFrac distances. PERMANOVA results testing differences in gut microbiome composition between pre-release and soft-release phases based on weighted UniFrac distances.

```
adonis2(formula = dist_wu_gut2 ~ phase + sex, data = meta_gut2, by = "margin")
         Df  SumOfSqs      R2      F Pr(>F)
phase     1 0.0002076 0.04450 0.6840  0.727
sex       1 0.0002073 0.04444 0.6831  0.621
Residual 14 0.0042492 0.91077
```

Table S20. PERMANOVA results for oral microbiomes using unweighted UniFrac distances. PERMANOVA results testing the effect of release candidacy on oral microbiome composition using unweighted UniFrac distances.

```
adonis2(formula = dist_uw_oral ~ status + sex, data = meta_oral, by = "margin")
         Df SumOfSqs      R2      F Pr(>F)
status    1   0.5805 0.09293 1.6379  0.007 **
sex       1   0.3585 0.05740 1.0116  0.394
Residual 15   5.3163 0.85104
Total    17   6.2469 1.00000
```

Table S21. PERMANOVA results for oral microbiomes using weighted UniFrac distances. PERMANOVA results testing the effect of release candidacy on oral microbiome composition using weighted UniFrac distances.

```
adonis2(formula = dist_wu_oral ~ status + sex, data = meta_oral, by = "margin")
         Df  SumOfSqs      R2      F Pr(>F)
status    1 0.0031337 0.15232 2.7832  0.002 **
sex       1 0.0007087 0.03445 0.6294  0.750
Residual 15 0.0168890 0.82090
Total    17 0.0205739 1.00000
```

---

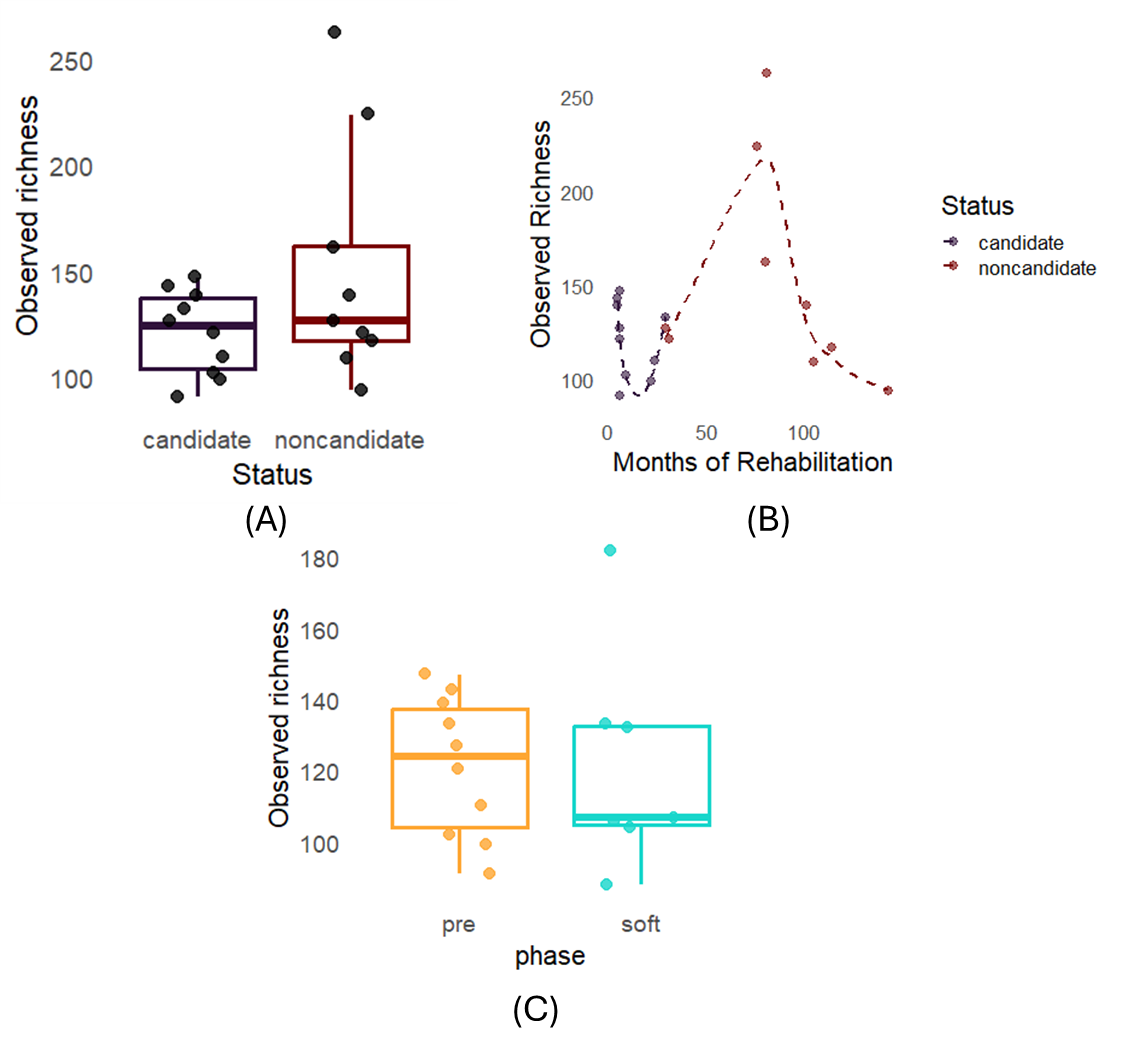

Figure S1. Gut microbiome alpha diversity (observed richness) across release status, rehabilitation duration, and release phase in Javan slow loris (*N. javanicus*). (A) observed ASV richness of gut microbiomes in candidate and non-candidate *Nycticebus javanicus*. (B) relationship between rehabilitation duration (months in captivity) and observed ASV richness of gut microbiomes for candidate and non-candidate individuals. Lines indicate locally weighted smoothing (LOESS) to visualize overall trends. (C) Observed ASV richness of gut microbiomes during pre-release and soft-release phases in candidate individuals. Points represent individual samples; boxplots show medians and interquartile ranges.

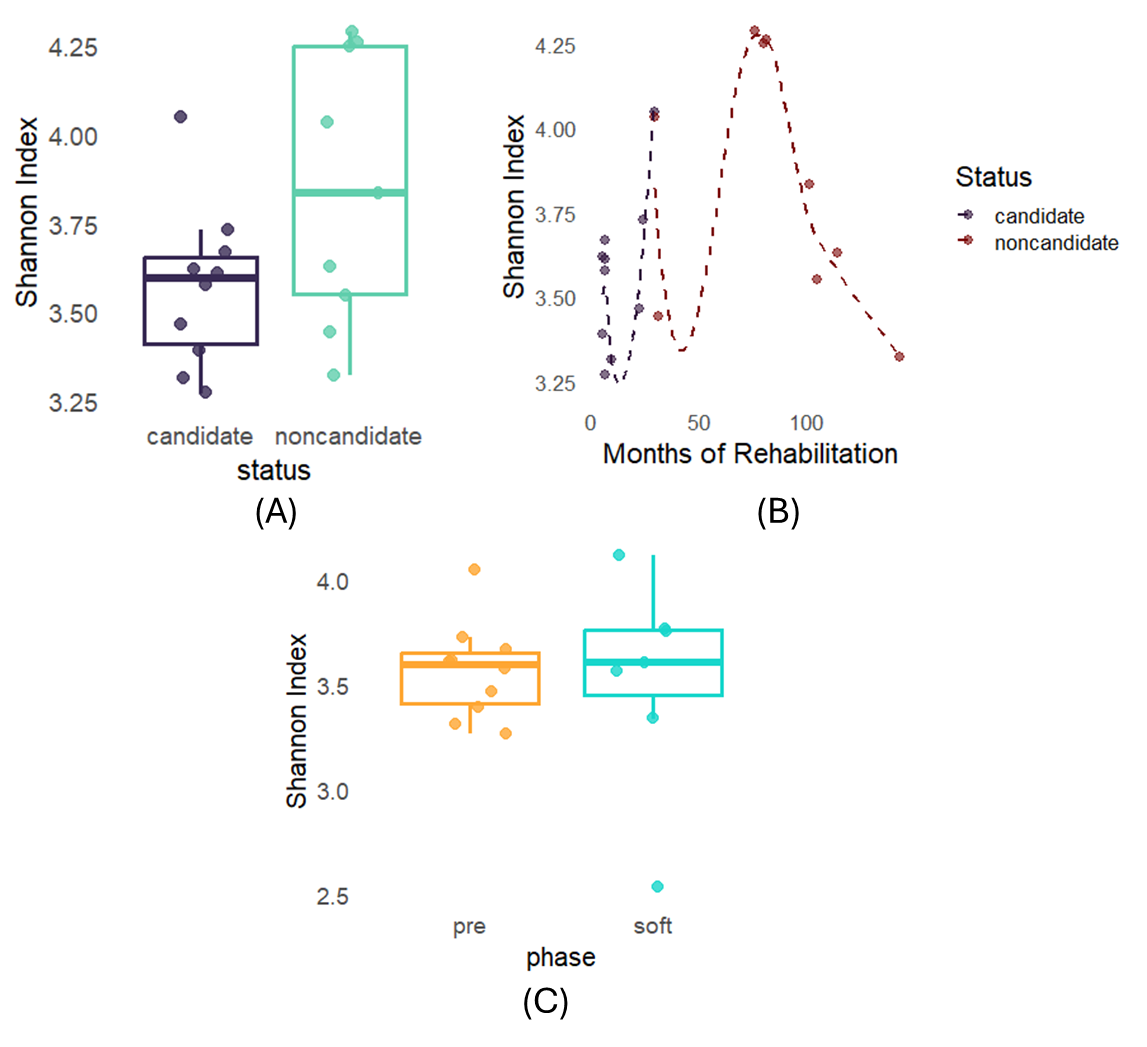

Figure S2. Gut microbiome alpha diversity (Shannon index) across release status, rehabilitation duration, and release phase in Javan slow loris (*N. javanicus*). (A) Shannon diversity of gut microbiomes in candidate and non-candidate *Nycticebus javanicus*. (B) Relationship between rehabilitation duration (months in captivity) and Shannon diversity of gut microbiomes for candidate and non-candidate individuals. Lines indicate locally weighted smoothing (LOESS) to visualize overall trends. (C) Shannon diversity of gut microbiomes during pre-release and soft-release phases in candidate individuals. Points represent individual samples; boxplots show medians and interquartile ranges.

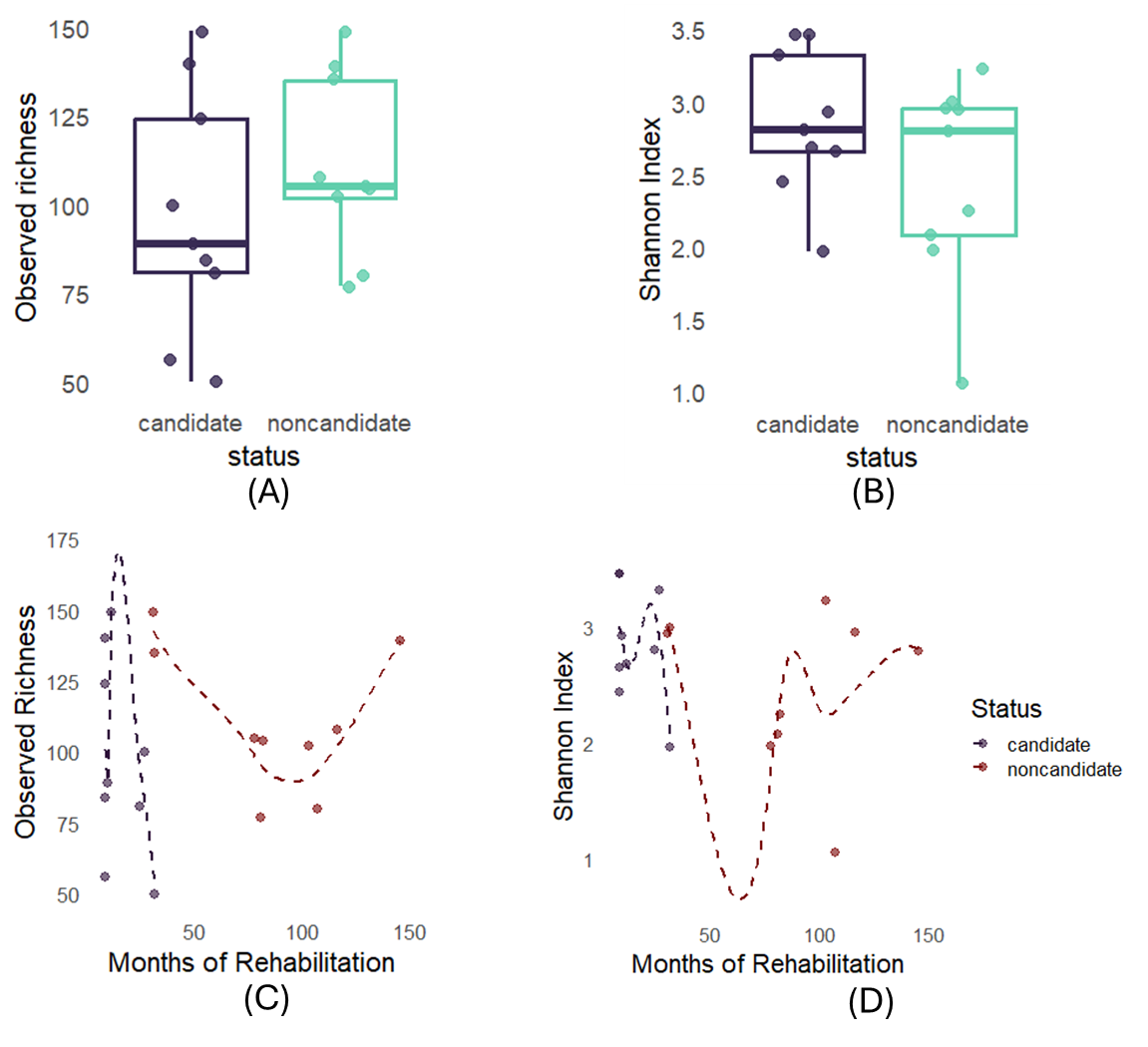

Figure S3. Oral microbiome alpha diversity of rehabilitating *Nycticebus javanicus*. (A) Observed ASV richness of oral microbiomes in candidate and non-candidate individuals. (B) Shannon diversity of oral microbiomes in candidate and non-candidate individuals. (C) Relationship between rehabilitation duration (months in captivity) and observed ASV richness of oral microbiomes. (D) Relationship between rehabilitation duration (months in captivity) and Shannon diversity of oral microbiomes. Points represent individual oral samples; boxplots show medians and interquartile ranges. Lines indicate locally weighted smoothing (LOESS) to visualize overall trends.

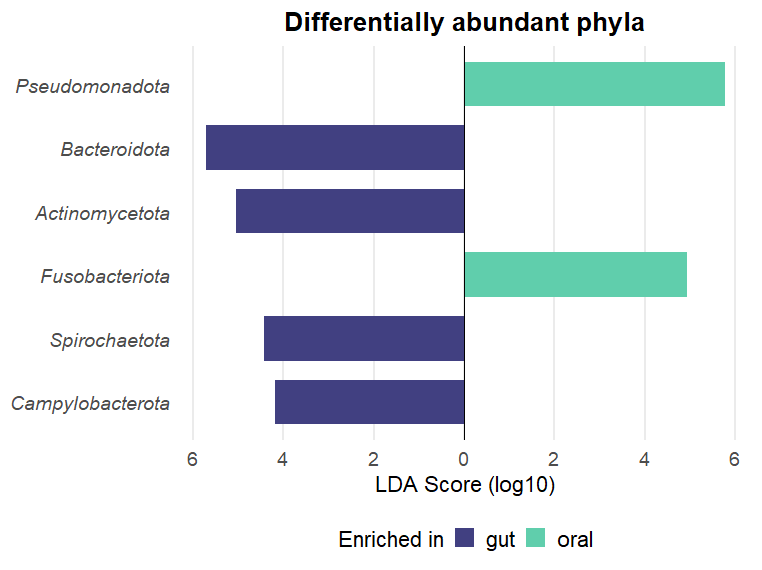

Figure S4. Differentially abundant bacterial phyla between gut and oral microbiomes of rehabilitating slow lorises (*Nycticebus javanicus)*. Linear discriminant analysis effect size (LEfSe) identifies bacterial phyla significantly enriched in gut (blue) or oral (green) microbiomes (LDA score ≥ 2.0, p < 0.05). Phyla enriched in the gut include Bacteroidota, Actinomycetota, Spirochaetota, and Campylobacterota, whereas the oral microbiome is characterized by enrichment of Pseudomonadota and Fusobacteriota. Bars represent log10-transformed LDA scores indicating the magnitude of differential enrichment.

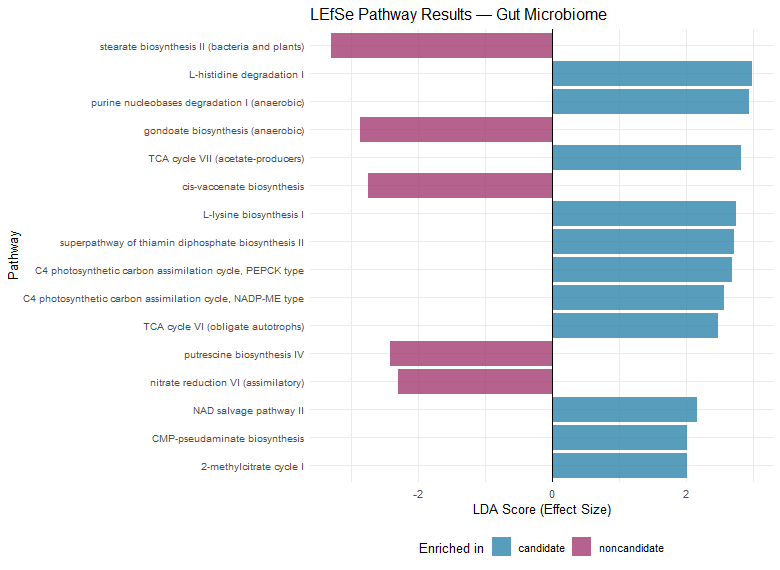

Figure S5. Differentially enriched predicted metabolic pathways in gut microbiomes of candidate and non-candidate slow loris (*Nycticebus javanicus)*. Predicted functional profiles of gut microbiomes were inferred using PICRUSt2 and compared between candidate and non-candidate individuals using linear discriminant analysis effect size (LEfSe). Bars indicate metabolic pathways that were significantly differentially enriched between groups (LDA score ≥ 2.0, p < 0.05). Pathways enriched in candidate individuals (blue) were primarily associated with central energy metabolism and biosynthetic functions, including multiple tricarboxylic acid (TCA) cycle variants, amino acid biosynthesis, vitamin and cofactor metabolism, and nucleotide metabolism. In contrast, non-candidate individuals (purple) showed enrichment of pathways related to lipid biosynthesis, polyamine metabolism (putrescine biosynthesis IV), and nitrate reduction, consistent with altered metabolic strategies under disturbed gut conditions. Bars represent effect sizes (LDA scores) indicating the magnitude of differential enrichment.

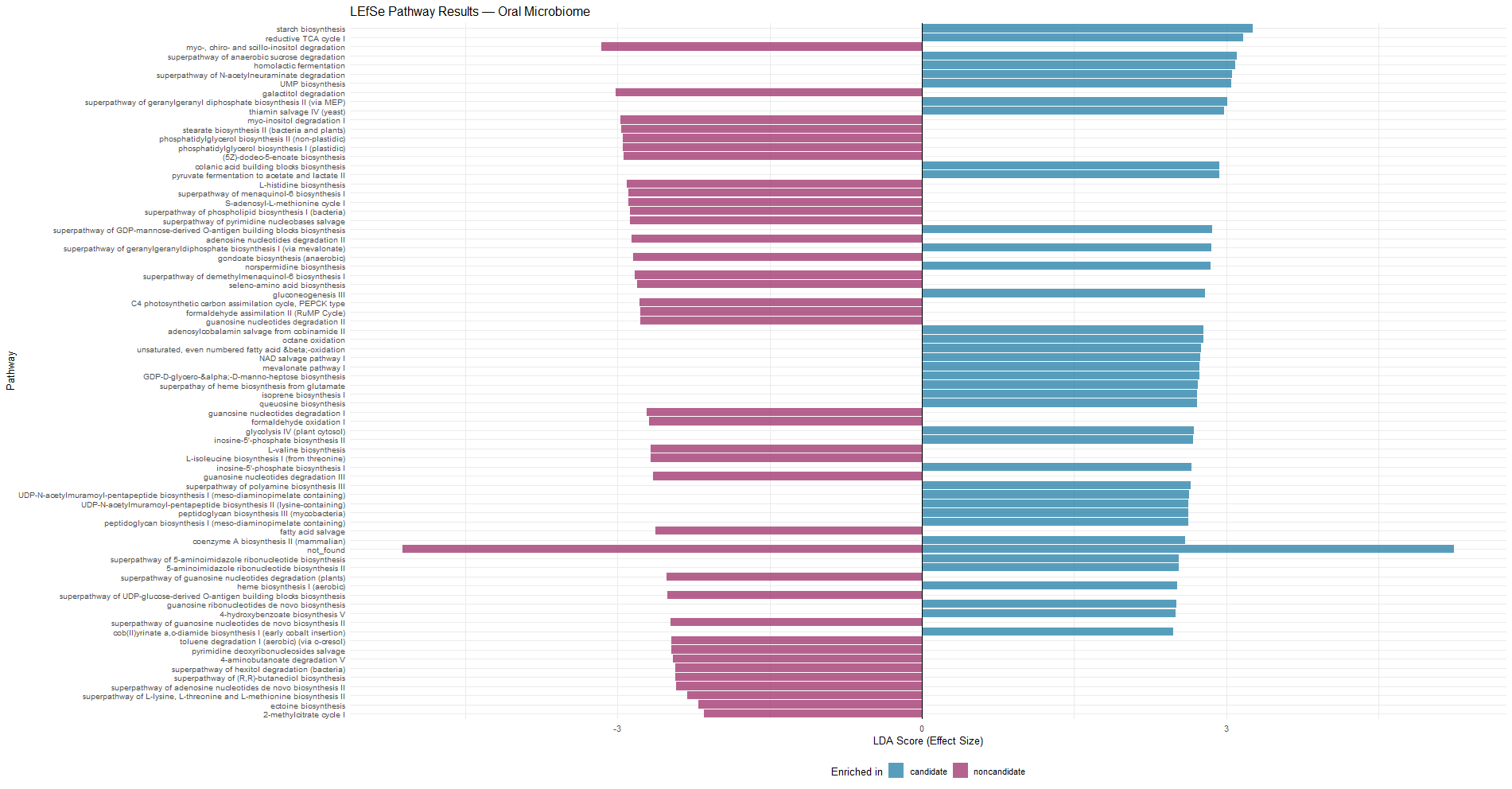

Figure S6. Differentially enriched predicted metabolic pathways in oral microbiomes of candidate and non-candidate slow loris (*Nycticebus javanicus*). Predicted functional profiles of oral microbiomes were inferred using PICRUSt2 and compared between candidate and non-candidate individuals using linear discriminant analysis effect size (LEfSe). Bars represent metabolic pathways that differed significantly between groups (LDA score ≥ 2.0, p < 0.05), with positive LDA scores indicating enrichment in candidate individuals (blue) and negative scores indicating enrichment in non-candidates (purple). Candidate oral microbiomes were enriched in biosynthetic and central metabolic pathways, including amino acid and nucleotide biosynthesis, vitamin and cofactor metabolism, and core energy metabolism. In contrast, non-candidate oral microbiomes showed enrichment of degradation- and recycling-associated pathways, including sugar alcohol degradation, aromatic compound degradation, nucleoside and nucleotide degradation, and lipid biosynthesis pathways. Bars indicate LDA effect sizes reflecting the magnitude of differential enrichment.
